# Noncanonical chromosomal-end-specific telomeric arrays in naturally telomerase-negative yeasts

**DOI:** 10.1101/2025.09.07.674783

**Authors:** Broňa Brejová, Viktória Hodorová, Hana Lichancová, Askar Gafurov, Dominik Bujna, Filip Brázdovič, Filip Červenák, Tomáš Petrík, Eva Hegedűsová, Michaela Forgáčová Jakúbková, Martina Neboháčová, Ľubomír Tomáška, Matthias Sipiczki, Tomáš Vinař, Jozef Nosek

## Abstract

In most eukaryotes, chromosomal DNA terminates with tandem repeats of a short G-rich motif, such as the canonical TTAGGG sequence. The arrays of telomeric repeats are maintained by telomerase or by alternative lengthening of telomeres (ALT). Here we report that nuclear chromosomes of several basidiomycetous yeasts classified into the order Microstromatales carry unusual telomeres. We demonstrate that instead of TTAGGG-like repeats these telomeres are composed of unique tandem arrays which are in most cases specific to a particular chromosomal end. In contrast to other basidiomycetes, the Microstromatales genomes lack orthologs coding for the telomerase catalytic subunit Est2 and a shelterin component Tpp1 indicating that noncanonical telomeric arrays are maintained by a telomerase-independent mechanism. We hypothesize that in a common ancestor of Microstromatales the loss of telomerase and Tpp1 was compensated by activation of an ALT mechanism, which promoted amplification of various motifs and formation of distinct telomeric arrays at most chromosomal ends.

## Introduction

Eukaryotic chromosomes terminate with specialized nucleoprotein structures termed telomeres that protect the genome integrity by preventing nucleolytic degradation and unwanted DNA repair activity (McEachern et al. 2000; de Lange, 2018). Telomeric DNA typically consists of arrays of a short G-rich motif repeated in tandem. In all major eukaryotic lineages, the motif is represented by the hexanucleotide TTAGGG. Its minor variations (*e*.*g*. TTAGG, TTTAGGG) were identified in protists, fungi, plants, and animals. Yet, noncanonical telomeric motifs occur in many species. Particularly puzzling variability was described in ascomycetous yeasts, whose telomeric sequences may be unusually long and/or irregular and may differ even between closely related species (Moyzis et al. 1988; Meyne et al. 1989; Gunišová et al., 2009; Fulnečková et al., 2013; Fajkus et al. 2016; Červenák et al. 2021; Peška et al. 2021). The length of telomeric arrays varies between organisms, from tens of base-pairs (bp) in ciliates (Klobutcher et al. 1981) up to above 150 thousand bp in mice (Kipling and Cooke, 1990) or even more, as has been reported in barley callus culture (Kilian et al. 1995). The very ends of chromosomes usually terminate with a 3’ single-stranded (ss) G-rich overhang of telomeric DNA (Klobutcher et al. 1981; Henderson and Blackburn, 1989) which has a propensity to fold into quadruplex (G4) structures (Tang et al. 2008). Mammalian telomeric DNA is bound by a six-subunit complex dubbed shelterin (Pot1–Tpp1–Tin2–Rap1–Trf1–Trf2) and a heterotrimeric protein complex CST (Ctc1–Stn1–Ten1), which mediate the chromosomal end protection and telomere length regulation (de Lange, 2018; Lim and Cech, 2021; Takai et al. 2024). Telomeric arrays are commonly synthesized by the enzyme telomerase reverse transcriptase (TERT), whose core components are the catalytic subunit Est2 and telomerase RNA (TR) providing both a structural scaffold for protein subunits and a template for the synthesis of telomeric repeats (Greider and Blackburn, 1987; Blackburn and Collins, 2011; Lue and Autexier; 2023). The telomerase provides a robust means of telomere maintenance. However, cells lacking its activity, including a significant subset of human cancers and immortalized cell lines, employ alternative lengthening of telomeres (ALT) mechanisms dependent on homologous recombination and DNA repair processes via break-induced replication (BIR) (McEachern and Haber, 2006; Cesare and Reddel, 2010; Roumelioti et al. 2016; Apte and Cooper, 2017; Hoang and O’Sullivan; 2020). The ALT pathways may operate both in telomerase-positive and telomerase-negative cells and represent a back-up mechanism which can be selectively induced in cells depleted of telomerase activity either due to a mutation or chemical inhibition. For example, surviving clones arising in a senescent population of mutant yeast *Saccharomyces cerevisiae* lacking functional telomerase amplify either subterminal Y’ elements (type I survivors requiring a strand invasion protein Rad51) or the telomeric repeats (type II survivors dependent on the MRX complex subunit Rad50 and DNA helicase Sgs1) eventually generating hybrid (type I/type II) arrays. Maintenance of these telomeres relies on recombinational protein Rad52 (Lundblad and Blackburn, 1993; Chen et al. 2001; Huang et al. 2001; Kockler et al. 2021). Yet, the telomerase-deficient mutants with long telomeric arrays may remain viable even in the absence of Rad52 employing the ‘inherited-long-telomere’ (ILT) pathway (Grandin and Charbonneau, 2009). In addition, yeast mutants defective for telomerase, Rad52, and exonuclease Exo1 generate survivors which stabilize their chromosomal termini by formation of palindromes (Maringele and Lydall, 2004). Analyses of *Schizosaccharomyces pombe* mutants revealed that the loss of telomerase can also be overcome by chromosome circularization (Nakamura et al. 1998) or the ‘heterochromatin amplification-mediated and telomerase-independent’ (HAATI) mechanism (Jain et al. 2010).

The ALT mechanisms represent the primary mode of telomere maintenance in eukaryotes that naturally lack telomerase. This can be exemplified by dipterans in which various noncanonical telomeres have evolved (Mason et al. 2016). Studies of chironomids (*e*.*g. Chironomus pallidivittatus*) and the mosquito *Anopheles gambiae* revealed that their chromosomal ends are composed of complex repeats elongated by a recombinational mechanism (Saiga and Edström, 1985; Nielsen and Edström, 1993; Roth et al. 1997). In contrast, the fruit fly *Drosophila melanogaster* maintains its chromosomal termini by transposition of non-LTR retrotransposons (Levis et al. 1993). Noncanonical telomeric repeats presumably maintained by an ALT mechanism were recently reported also in several nematode species from the genus *Meloidogyne* (Mota et al. 2024) and *Strongyloides stercoralis* (Chung et al. 2025). Interestingly, the first case of a vertebrate naturally lacking the telomerase has been reported in highly regenerative newt *Pleurodeles waltl*, whose chromosomes terminate with interspersed variant repeats which are maintained by an ALT mechanism (Yu et al. 2022).

In this study we show that *Jaminaea angkorensis*, a basidiomycetous yeast taxonomically classified into the order Microstromatales (class Exobasidiomycetes, subphylum Ustilaginomycotina), lacks canonical TTAGGG-like telomeric arrays maintained by telomerase which are typical for basidiomycetes (Guzmán and Sánchez, 1994; Sanpedro-Luna et al. 2023; Sanpedro Luna and Sánchez Alonso, 2025), including the model species, corn smut *Mycosarcoma maydis* (formerly *Ustilago maydis*; McTaggart et al. 2016). By combination of nuclease mapping, long read sequencing and bioinformatic analysis, we demonstrate that 20 chromosome-sized contigs in the genome assembly of *J. angkorensis* terminate with noncanonical telomeric sequences. The very ends in 36 out of 40 chromosomal termini are composed of distinct motifs varying both in size and nucleotide composition. Some chromosomes terminate with complex arrays, composed of distinct proximal and distal repeats. For comparative purposes, we generated the chromosome-level genome assemblies of additional three Microstromatales species, *Jaminaea pallidilutea, Parajaminaea (Jaminaea) phylloscopi*, and *Sympodiomycopsis kandeliae*. These species also lack TTAGGG-like telomeres, and their chromosomes terminate with various noncanonical repeats distinct from those identified in *J. angkorensis*. We also show that all investigated species lack the genes for the catalytic subunit of telomerase Est2/TERT and a component of the shelterin complex Tpp1, which forms a structured interface with N-terminal domain of Est2 and is crucial for chromosomal end protection as well as telomerase recruitment to telomeric DNA (Liu et al. 2022; Sekne et al. 2022). Except for Est2 and Tpp1, the orthologs coding for other key proteins involved in telomere maintenance are encoded in the Microstromatales genomes. As both the telomerase and TTAGGG-like telomeric repeats commonly occur in basidiomycetes, we hypothesize that noncanonical telomeres in Microstromatales emerged by activation of ALT which compensated for the loss of telomerase-dependent telomere synthesis and the complete erosion of ancestral telomeric arrays.

## Results

### Nuclear chromosomes of *J. angkorensis* terminate with noncanonical, chromosomal-end-specific telomeric arrays

To investigate the genetic make-up of *J. angkorensis*, we first assembled its genome sequence using long reads generated by nanopore technology on a MinION device (Oxford Nanopore Technologies) and polished it using short Illumina reads (**Supplementary Table S1**). The resulting genome assembly (20.75 Mbp, 60.3% G+C) consists of 20 nuclear contigs with sizes ranging from 0.1 to 2.9 Mbp and a 30 kbp long contig representing a circular mitochondrial DNA (**Table 1**; **Supplementary Table S1**). The nuclear contigs correspond to the profile of the electrophoretic karyotype determined by pulsed-field gel electrophoresis (PFGE), although several chromosomes with similar sizes co-migrate as groups and could not be resolved into individual bands (**Supplementary Figure S1**). The PFGE results indicate that the nuclear contigs likely represent the whole chromosomes. Surprisingly, these contigs lack the telomeric repeats commonly occurring in basidiomycetes (*e*.*g*. Guzmán and Sánchez, 1994). Instead, we found that they terminate with tandem arrays of various noncanonical motifs. Since our attempts to identify TTAGGG-like repeats both in the genome assembly and the raw sequencing reads failed, we decided to map the sequences at the chromosomal termini experimentally. Our approach was modified from the original method developed by Peška et al. (2015, 2017). It is based on the depletion of telomeric repeats in sequencing reads generated by next-generation sequencing (NGS) of high molecular weight (HMW) genomic DNA digested with BAL-31 nuclease. This enzyme specifically shortens the ends of linear DNA molecules, so, compared with the untreated control, the nuclease-treated DNA samples are depleted of the terminal sequences. Specifically, we isolated the genomic DNA from *J. angkorensis* and digested it with BAL-31 nuclease. We performed two experiments, each with two different conditions for the nuclease reaction. In the first experiment, we used ~0.2 U of BAL-31 per μg of DNA and the samples were digested for 15 and 30 min. In the second experiment, the enzymatic digestion was performed for 30 min with ~0.2 U and ~0.6 U of BAL-31 per μg of DNA. The DNA fragments from the digested and control samples were then barcoded and sequenced using a tagmentation kit on a nanopore sequencer. In the obtained nanopore reads (**Supplementary Table S1**), we counted the occurrences for each 21-mer present in the genome assembly. We then highlighted 1 kbp windows that were more than 50% covered by 21-mers that were significantly depleted in the treated sample, compared with the control (**Figure 1**). The sequence near each contig end was depleted in at least one experiment, and as expected, the samples with a higher dose of BAL-31 or longer treatment time generally show more depleted terminal sequences. Only a single region distant from chromosome ends was depleted in one of the experiments. This region is located on chromosome 4 and contains long arrays of ribosomal DNA (rDNA), which are known to be unstable and eventually can generate long linear DNA fragments (Zylstra et al. 2023). The BAL-31 experiments confirmed that the chromosomal ends of all 20 nuclear chromosomes of *J. angkorensis* terminate with tandem arrays of various noncanonical motifs. Overall, we identified 36 unique terminal repeat motifs. Their sizes and G+C content vary from 52 to 178 bp and 41.4 to 66.3%, respectively (**Table 1**; **Supplementary Table S2**; **Supplementary Table S3**). While most chromosomal ends terminate with an array composed of a single distinct motif, chromosome 20 contains arrays of the same 130 bp long repeat at both ends. In addition, several chromosomal ends are more complex and consist of the proximal and distal arrays. The right ends of chromosomes 2, 8, and 13 possess distal arrays of the same 52 bp long motif, yet these termini differ in the proximal arrays consisting of motifs with the sizes of 80, 126, and 72 bp, respectively. Similarly, a 178 bp long motif occurs at the distal arrays of the right telomere of chromosomes 4 and 18, whose proximal arrays are composed of 73 and 63 bp long motifs, respectively. Taken together, the terminal regions of 19 out of 20 chromosomes appear to be unique, represented either by an array of a single distinct telomeric repeat or by a complex of proximal and distal arrays as described above.

**Table 1.**
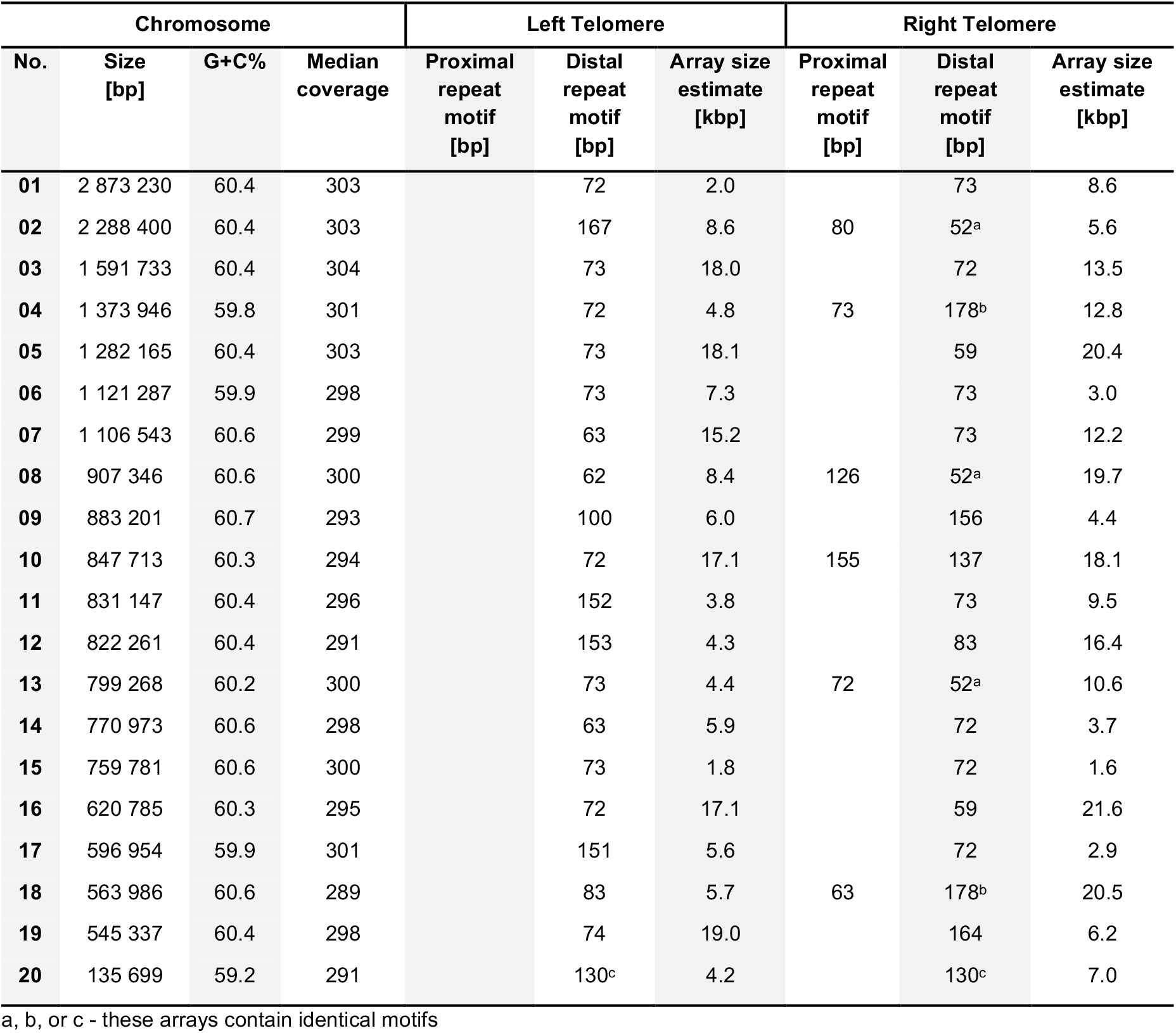
Characteristics of telomeric arrays in *J. angkorensis*.

**Figure 1.**
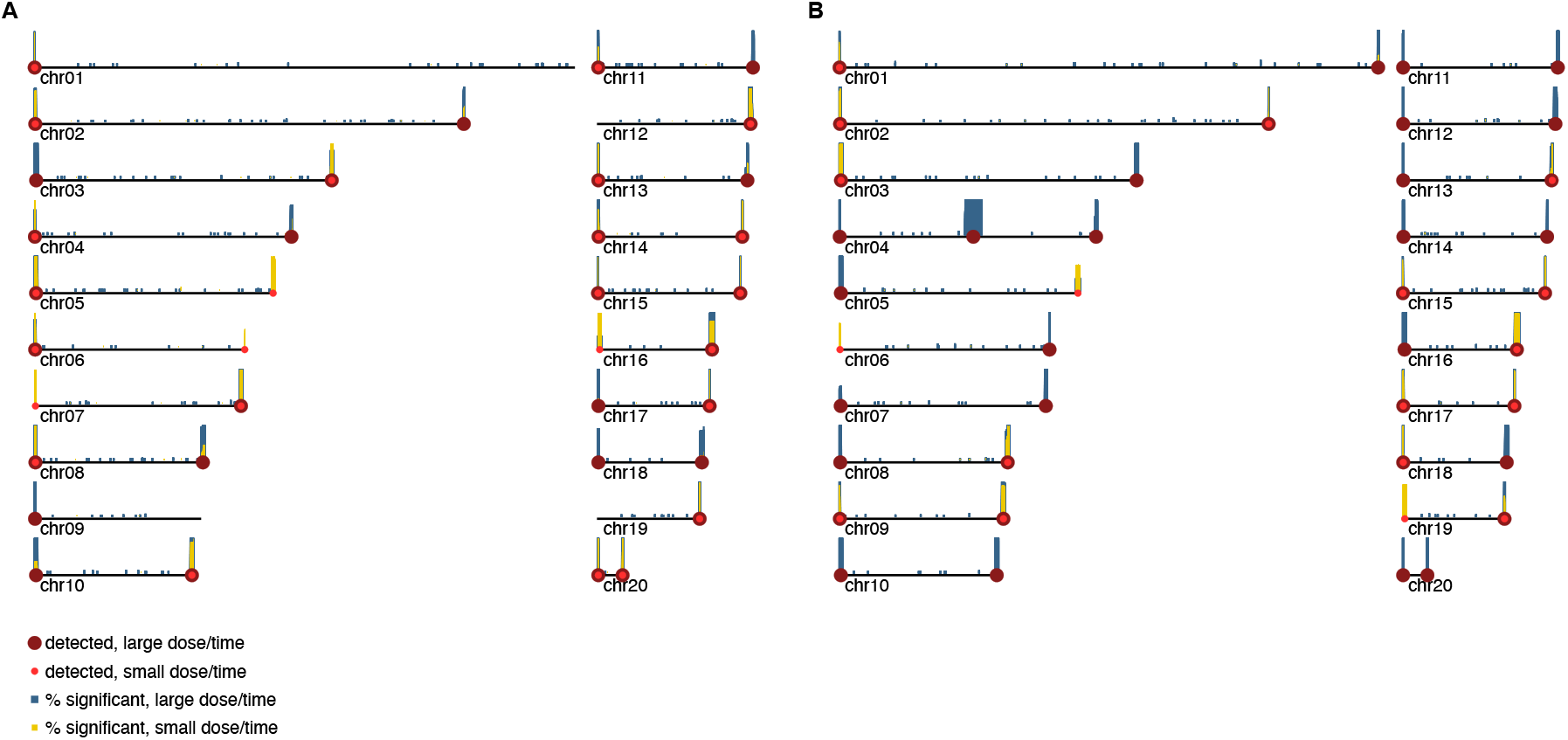
Genomic regions with decreased nanopore sequencing coverage after BAL-31 treatment. Yellow and blue bars show the coverage of 1 kbp windows by k-mers significantly depleted by the BAL-31 treatment, red circles highlight the regions where this coverage exceeds 50%. (A) The first experiment, with ~0.2 U of BAL-31 per μg of DNA and the treatment times 15 and 30 min. (B) The second experiment, with ~0.2 U and ~0.6 U of BAL-31 per μg of DNA and the treatment time 30 minutes. In both plots, darker colors and larger circles are used for longer treatment or a larger enzyme dose.

To further confirm the identified telomeric motifs, we chose the sequences present at the termini of chromosomes 15 and 20 and tested their sensitivity to BAL-31 nuclease using a conventional terminal restriction enzyme fragment (TRF) analysis. Our results show that these sequences are sensitive to increasing amounts of the nuclease, thus confirming their location at the chromosomal ends (**Supplementary Figure S2**). Moreover, the subsequent analysis of nanopore reads revealed that the sizes of the repetitive arrays at the ends of individual chromosomes differ by more than tenfold (from ~1.6 to ~21.6 kbp; **Figure 2**; **Table 1**; **Supplementary Table S2**). The variable TRF lengths were also demonstrated by Southern blot hybridizations with oligonucleotide probes derived from telomeric repeats of chromosomes 1, 3, 15, and 20 (**Supplementary Figure S3**).

**Figure 2.**
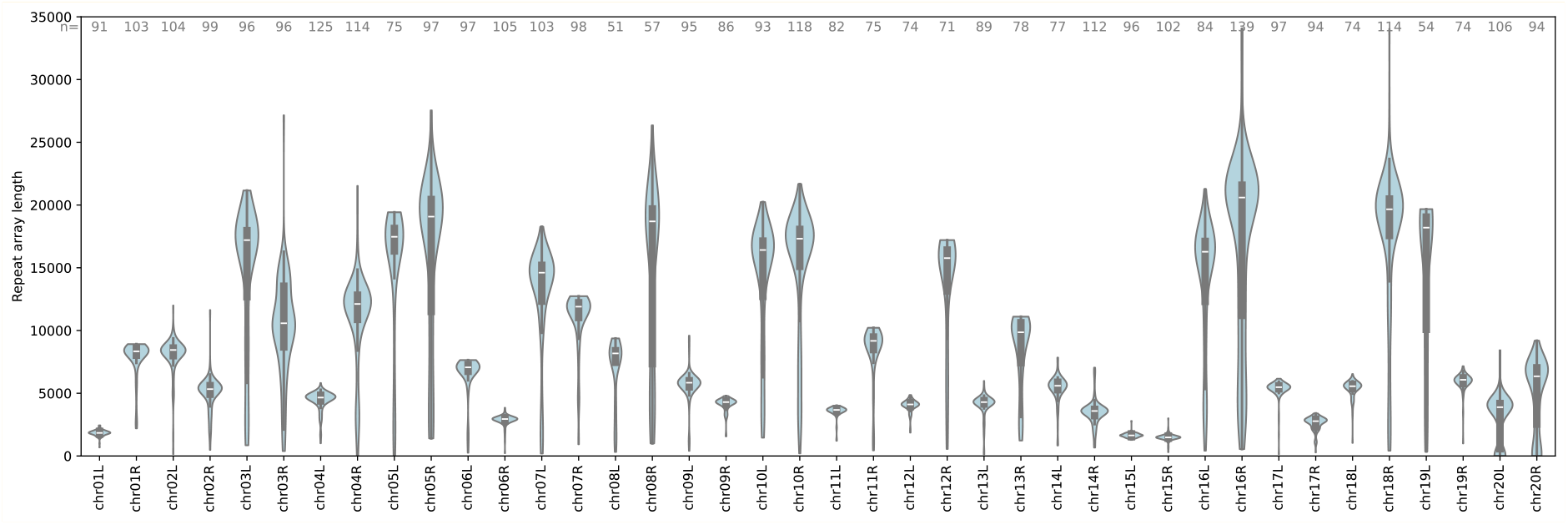
Violin plots of the lengths of telomere motif arrays in individual nanopore sequencing reads for each chromosomal end in *J. angkorensis*. Both distal and proximal motifs are considered in the length. Only reads spanning a unique sequence region closest to chromosome end and longer than 30 kbp were considered. Nonetheless some reads may represent fragments not reaching the actual chromosome length. The numbers at the top of the plot show the read count used in each violin plot.

To investigate whether noncanonical telomeres of *J. angkorensis* terminate with a 3’ ss overhang, which is considered a conserved feature of telomeres in most eukaryotes (Henderson and Blackburn, 1989), we digested its genomic DNA with exonuclease I (ExoI), RecJ_f_, and mung bean nuclease (MBN). These enzymes exhibit 3’ to 5’ (ExoI) and 5’ to 3’ (RecJ_f_) ss-specific exonuclease and ss-specific endonuclease (MBN) activity. Digested DNA samples, together with undigested native and denatured DNA controls, were blotted and hybridized with radiolabeled ss-oligonucleotides derived from telomeric motifs present at both ends of chromosome 15 (G-rich probe) as well as their complementary counterparts (C-rich probe). Our results revealed the telomeric ssDNA regions detectable in native controls, which are almost completely degraded by MBN. In contrast to MBN, the treatment with the exonucleases ExoI and RecJ_f_ did not remove the signal detected in native controls. Moreover, in contrast to canonical telomeres, the ssDNA regions were detected by both the G-rich and C-rich probes (**Supplementary Figure S4**). These results indicate that rather than overhangs at the very ends of *J. angkorensis* chromosomes, the identified ssDNA regions may represent gaps within the internal parts of the telomeric arrays. Such gaps could result from incomplete DNA synthesis and/or unrepaired recombination events occurring within the telomeric arrays.

### Noncanonical telomeres are common in the order Microstromatales

Surprising molecular organization of *J. angkorensis* telomeres prompted us to analyze the chromosomal termini of three additional species *J. pallidilutea, P. phylloscopi*, and *S. kandeliae*, which also belong to the order Microstromatales. As in the case of *J. angkorensis*, we assembled their genome sequences using nanopore and Illumina reads (**Supplementary Table S1**). The nuclear genome assemblies of these species contain 16, 3, and 6 contigs, respectively (**Supplementary Table S1**), corresponding to their electrophoretic karyotypes (**Supplementary Figure S1**), and therefore they likely represent complete chromosomes. The four Microstromatales genomes differ by the overall size, the G+C content and the number of chromosomes, which vary from 17.2 Mbp (*J. pallidilutea*) to 20.7 Mbp (*J. angkorensis*), from 49.8% (*S. kandeliae*) to 60.3% (*J. angkorensis*), and from 3 (*P. phylloscopi*) to 20 (*J. angkorensis*), respectively. The chromosome sizes in these species range from ~ 0.1 to 9.9 Mbp (**Supplementary Table S1, Supplementary Figure S1**).

Similarly to *J. angkorensis*, the chromosomal contigs of these species lack TTAGGG-like repeats and their ends are composed of arrays of a single noncanonical motif, which is in most cases chromosomal-end-specific, or they contain a combination of proximal and distal repeats (**Supplementary Table S2, Supplementary Table S3**). In the distal arrays of *J. pallidilutea, P. phylloscopi*, and *S. kandeliae*, we identified 18, 5, and 7 unique motifs, respectively, with sizes ranging from 41 to 100 bp, 101 to 132 bp, and 55 to 122 bp. The subsequent analysis of nanopore sequencing reads indicated that also the lengths of individual telomeric arrays are highly variable (**Supplementary Figure S5**). We found that in each species, some motifs are repeated at different chromosomal ends. To characterize the similarities between individual motifs more broadly, we compared their 7-mer compositions (**Figure 3**). As a result, we found out that some groups of motifs within a single species are highly similar (7-mer containment distance less than 0.1, see Methods). The similarity of motifs among different species is limited (closest pairs are at a distance of 0.71, involving a group of three *J. pallidilutea* and three *S. kandeliae* motifs) and even within a single species, we see multiple groups of very distant motifs. Furthermore, no interspecific homology was found by Blast with E-value threshold 10^-5^. This suggests that telomeric motifs do not share a single common evolutionary origin or they mutate at a very high rate. Nonetheless, we observed several large conserved chromosomal blocks in the whole-genome alignments (**Supplementary Figure S6**).

**Figure 3.**
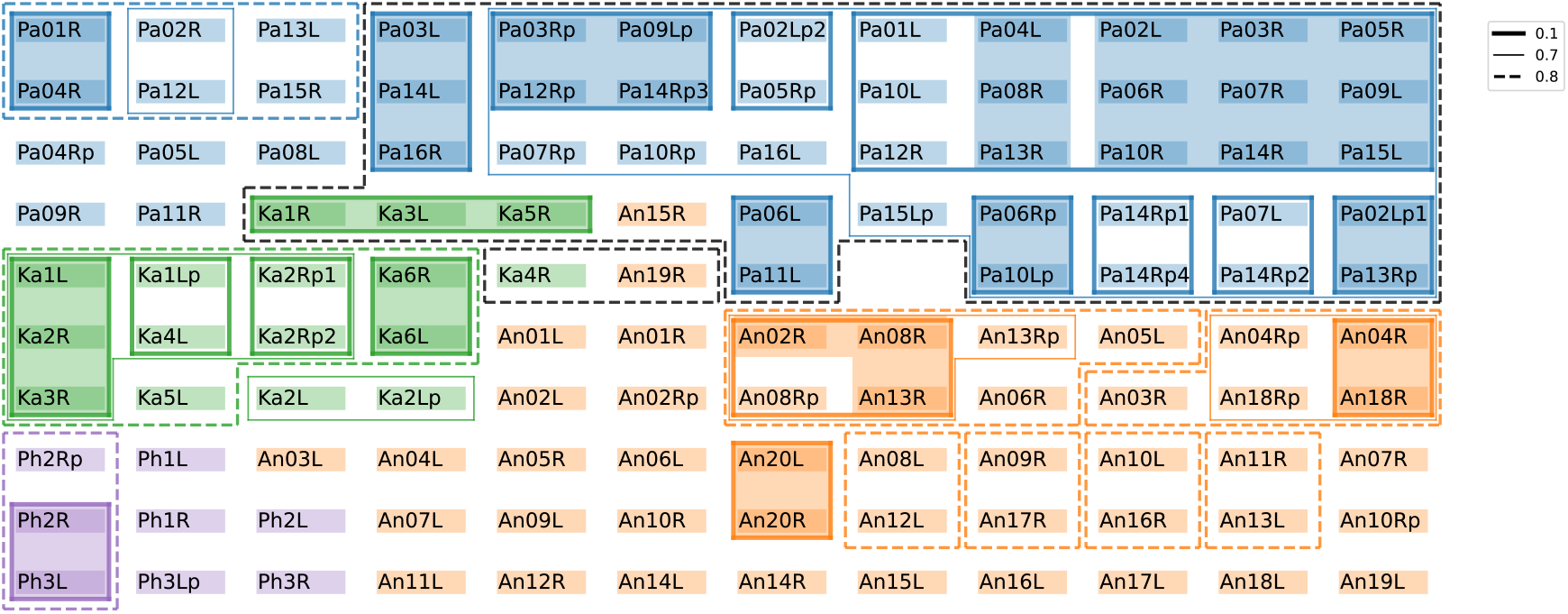
Single-linkage clustering applied to telomere motifs from all four studied species with 7-mer containment distance. (see Methods). Each motif is shown as a rectangle labeled with a code consisting of the first 2 letters of the species name, chromosome number, left (L) or right (R) chromosomal end and letter p denoting proximal pattern. For example, Pa01R is the distal motif at the right end of chromosome 01 of *J. pallidilutea*. Groups of motifs clustered at three different distance thresholds are enclosed by polygons of different line styles. Identical motifs are shown by a shaded background.

### Inventory of proteins involved in telomere maintenance

The diversity of noncanonical telomeric arrays identified in all four Microstromatales species also raises the question on the mode of telomere maintenance. To provide an insight into its molecular nature, we investigated the inventory of proteins potentially associated with telomeres or mediating telomeric functions. We annotated protein-coding genes in the genome assemblies using Augustus gene prediction software (Stanke et al. 2006) supported by RNA-seq, followed by manual curation of selected genes. In total, we identified 8263, 6317, 6583, and 7053 protein-coding genes in *J. angkorensis, J. pallidilutea, P. phylloscopi*, and *S. kandeliae*, respectively. We searched the predicted proteomes as well as the genome sequences using the BlastP and tBlastN tools (Altschul et al. 1997), and the queries derived from amino acid sequences of twenty-eight proteins from *M. maydis*, which represents the best characterized basidiomycete model organism in telomere biology and belongs to the same subphylum Ustilaginomycotina as the order Microstromatales. The queries included protein subunits of telomerase (Est1, TERT/Est2), components of the shelterin complex (Pot1, Rap1, Tpp1, Trf1/Tay1, Trf2), the CST complex (Stn1, Ten1), the Ku70/80 complex (Ku70, Ku80), the MRN/MRX complex (Mre11, Rad50, Nbs1), ATR/ATM kinases (Mec1, Tel1), DNA helicases (Blm/Sgs1, Dna2, Pif1, Rad54, Srs2), recombinational protein Rad52 and the members of Rad51 family (Rad51, Rad57/Rec2), a homolog of BRCA2 Brh2, exonuclease Exo1, a flap endonuclease Rad27/Fen1, and the ASTRA complex subunit (Tel2). The predicted proteomes were also searched using hhmsearch and hidden Markov model (HMM) profiles of the corresponding protein domains (Eddy, 2011). Note that homologs of the shelterin component Tin2 and Ctc1 subunit of the CST complex were not identified in *M. maydis* (Yu et al. 2013) and the searches with human protein queries or the corresponding protein domains did not reveal any significant hit in any examined species. For comparative purposes, we also performed these searches in two additional species - *Jaminaea rosea* and *Pseudomicrostroma glucosiphilum* from the order Microstromatales, as well as in five species (*Acaromyces ingoldii, Ceraceosorus guamensis, Meira miltonrushii, Tilletiaria anomala, Tilletiopsis washingtonensis*) classified into other orders of the class Exobasidiomycetes that possess canonical telomeric repeats. Our searches revealed that except for Est2 and Tpp1, the Microstromatales genomes code for the homologs of all searched protein queries (**Figure 4**). The absence of the TERT catalytic subunit Est2 and the shelterin component Tpp1, which is involved in telomerase recruitment to telomeres, indicates that noncanonical telomeres in Microstromatales are maintained by a telomerase-independent mechanism. Moreover, the presence of homologs of other telomeric proteins including those involved in DNA recombination and repair indicates that *J. angkorensis* and other closely related species contain the inventory of molecular tools required for processes involved in telomere maintenance in telomerase-negative ALT cells. The absence of TTAGGG-like repeats further points to similarity with the type I survivors emerging in cultures of *M. maydis* mutants lacking telomerase, that carry telomeres with amplified subterminal repeats and require the activities of Rad51 and Brh2 (Yu et al. 2017). We found that Est2 and Tpp1 homologs are missing also in the proteomes of *J. rosea* and *P. glucosiphilum*, whose available genome assemblies lack the TTAGGG arrays. Therefore, we presume that these two species also possess the noncanonical telomeric repeats, although their sequences are yet to be determined.

**Figure 4.**
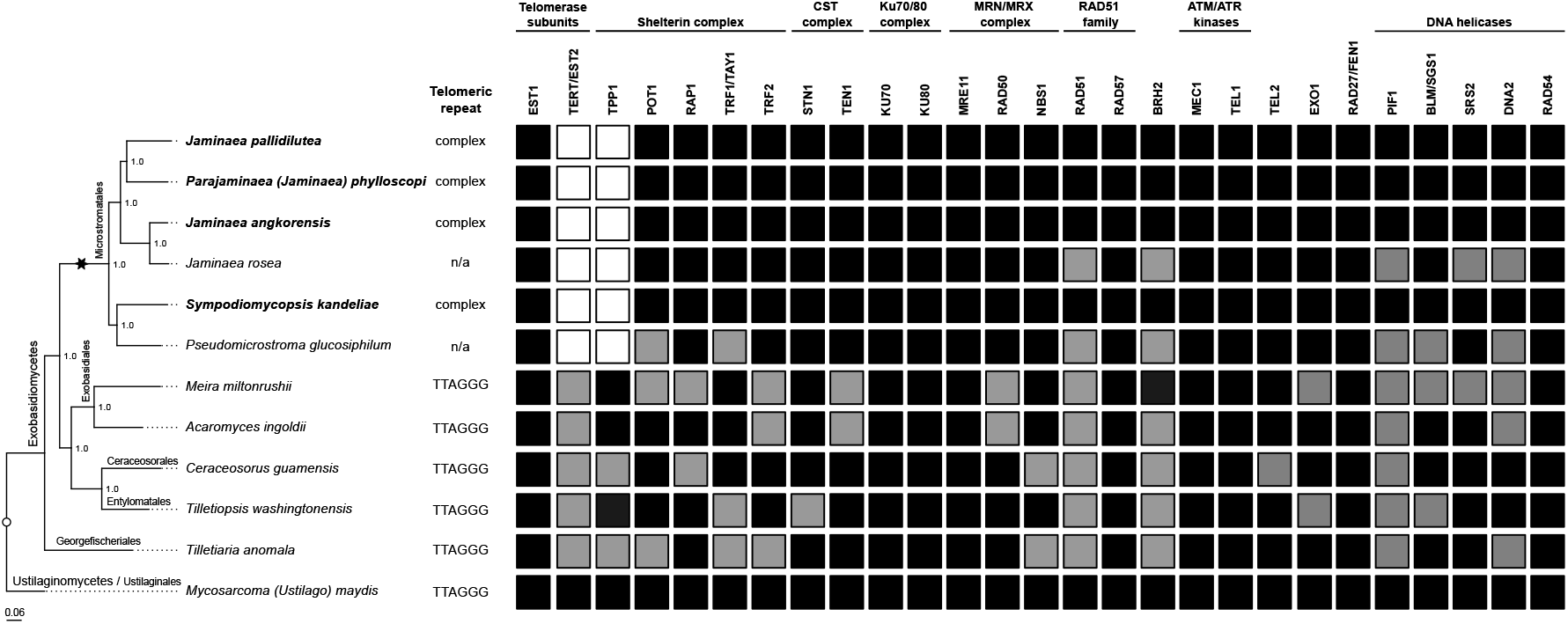
Phylogenetic tree of *J. angkorensis* and related Microstromatales species whose chromosomes terminate with noncanonical repeats. The phylogenetic tree was built from 1,764 single-copy orthologs present in at least 50% of the reference genome assemblies of indicated species; *M. maydis* was used as an outgroup (see Supplementary Methods for details). Analyses of the genome sequences revealed that species classified in the order Microstromatales (lineage is indicated by an asterisk) lack TTAGGG telomere motif as well as orthologs coding for the catalytic subunit of telomerase Est2 and a shelterin component Tpp1. Black and grey squares represent the presence of orthologs of *M. maydis* proteins identified in the predicted proteomes or the genome assemblies by BlastP and tBlastN searches, respectively. The absence of an ortholog is indicated by an empty square; n/a - telomeric repeats are missing in published genome assemblies of *J. rosea* and *P. glucosiphilum*.

## Discussion

Although ascomycetous yeasts *S. cerevisiae* and *S. pombe* have proven to be excellent model organisms in telomere biology, telomeres in basidiomycetes appear more similar to the ends of mammalian chromosomes. This can be exemplified by *M. maydis*, whose telomeric arrays are composed of canonical, human-like motif TTAGGG (Guzmán and Sánchez, 1994). Other similarities with human cells include the composition of the shelterin complex and the role of *M. maydis* orthologs of recombination protein Rad51, a BRCA2 homolog Brh2, a DNA helicase Blm as well as the Ku70/80 and MRN/MRX complexes in telomere maintenance (Yu et al. 2013; Yu et al. 2015; Yu et al. 2018; Yu et al. 2020; Zahid et al. 2022; Syed et al. 2024).

In this study, we show that basidiomycetous yeasts classified into the order Microstromatales are naturally occurring telomerase-deficient organisms. These species lack the orthologs encoding the catalytic subunit of telomerase and the shelterin component Tpp1, as well as canonical, TTAGGG-like telomeric repeats typical for basidiomycetes. We found that chromosome-sized contigs in the *J. angkorensis* nuclear genome assembly terminate with a plethora of noncanonical arrays that, in most cases, are specific to a particular chromosomal end. The sensitivity of these arrays to BAL-31 nuclease (**Figure 1, Supplementary Figure S2**) indicates that these repeats represent genuine telomeric DNA. This conclusion is further supported by the analysis of chromosome-level genome assemblies of the other three species, *J. pallidilutea, P. phylloscopi*, and *S. kandeliae*, whose chromosomal termini also contain various noncanonical repeats. Similar to *J. angkorensis*, all three species lack homologs of the *EST2* and *TPP1* genes. We also found that these two genes as well as the canonical telomeric repeats are missing in the published genome sequences of additional two species from this phylogenetic lineage, *J. rosea* and *P. glucosiphilum* (Kijpornyongpan et al. 2018). Although some contigs in the genome assemblies of these two species terminate with tandem repeats of relatively long motifs, the nature of their telomeres remains unknown as these assemblies have not yet been finalized to the level of complete chromosomes. Nevertheless, our results indicate that the absence of Est2, Tpp1, and TTAGGG-like telomeric arrays is a common feature of the order Microstromatales, making these yeasts particularly suitable for the investigation of telomerase-independent mechanisms of telomere maintenance and their evolutionary origin. The Microstromatales species belong to the class Exobasidiomycetes, whose other lineages possess Est2, Tpp1 as well as the canonical telomeric repeats (**Figure 4**). It is therefore likely that the ancestor of Microstromatales contained TTAGGG-like telomeric arrays, and their complete erasure was a direct consequence of telomere shortening in the absence of telomerase activity. Interestingly, except the rDNA repeats, the proximal and distal telomeric repeats represent the most prominent tandem repeat arrays in the examined Microstromatales genomes (**Supplementary Figure S7**).

The sudden change of telomeric DNA sequence from TTAGGG-like repeats to arrays of noncanonical motifs may provide a burden on the stability of nucleoprotein complexes at the chromosomal ends. Conversely, structural alterations of these complexes may facilitate the sequence alterations. For example, the diversification of telomeric DNA motifs in distinct lineages of ascomycetous yeasts was accompanied by recruitment of different telomeric DNA-binding proteins (TBPs), such as Rap1, Tay1, and Taz1, whose orthologs are loosely conserved indicating they underwent a rapid evolution. In addition, experimental studies of DNA-binding properties revealed that these proteins recognize a broader spectrum of target motifs which opens a space for changes of telomeric sequences (Steinberg-Neifach and Lue, 2015; Sepšiová et al. 2016; Červenák et al. 2021; Lue, 2021). This can be exemplified by Tay1 from the ascomycetous yeast *Yarrowia lipolytica*, which binds both *Y. lipolytica*-specific and human-like telomeric motifs (Višacká et al. 2012). Rapid diversification of the Tay1 protein family in fungi further underscores possible alterations of its DNA-binding properties (Lue, 2021). Similarly, limited DNA-binding specificity of conserved TBP CEH-37 to various telomeric motifs is thought to underlie the diversification of telomeric repeat motifs in nematodes (Song et al. 2025).

In the genome sequences of Microstromatales, we identified an ortholog of Trf1/Tay1 along with three other components of the shelterin complex (Trf2, Pot1, and Rap1), although these yeasts lack Tpp1 (**Figure 4**). In mammalian cells, Tpp1 does not bind directly to the telomeric DNA, it forms a stable heterodimer with ssDNA-binding protein Pot1 (Chen et al. 2017; Rice et al. 2017), interacts with Tin2 (Takai et al. 2011), recruits telomerase to telomeric DNA and stimulates its processivity (Zaug et al. 2010; Sandhu et al. 2021). Mice with Tpp1 null mutation die perinatally and their cells exhibit accelerated telomere shortening as well as increased frequency of chromosome end fusions and fragility demonstrating that Tpp1 also prevents the activation of DNA damage response (DDR) and maintains the stability of chromosomal ends (Tejera et al. 2010). Notably, the selective degradation of Tpp1 during HSV-1 virus infection results in a rapid loss of telomeric repeats, dissociation of shelterin components from telomeres, DDR activation, and promotion of the viral DNA replication, presumably by recruiting telomere associated proteins to the virus replication compartments (Deng et al. 2014). The loss of Tpp1 thus results in uncapped chromosomes which tend to undergo various rearrangements eventually leading to karyotype alterations. It is therefore of note that the number of chromosomes in the examined Microstromatales species vary from 3 (*P. phylloscopi*) up to 20 (*J. angkorensis*) and the interspecies comparison revealed that they underwent numerous rearrangements (**Supplementary Figure S6**). It is also conceivable that concerted loss of Est2 and Tpp1 in an ancestor of Microstromatales led to the complete removal of ancestral TTAGGG-like repeats along with remodeling of telomeric nucleoprotein complexes that could be prerequisites for the emergence of noncanonical arrays which in turn stabilized the chromosomal ends.

While most telomeric arrays in the studied Microstromatales species consist of a unique motif, the ends of several chromosomes are complex and contain combinations of proximal and distal repeat arrays. In a subset of telomeres, the identified distal arrays are composed of the same motif which indicates that the motif could expand to the ends of multiple chromosomes. This points to two intriguing evolutionary scenarios. In the first, the proximal arrays may represent remnants of former telomeric repeats that were capped by arrays of a new motif. The occurrence of several types of proximal repeats on some chromosomal ends of *J. pallidilutea* (*i*.*e*. the left and right telomere of chromosome 2 and 14, respectively) and *S. kandeliae* (*i*.*e*. the right telomere of chromosome 2) is suggestive of multiple rounds of such replacements. This idea is also supported by *P. phylloscopi* chromosome 2, where the 101 bp long motifs present at the distal array of the left telomere and the proximal array of the right telomere are highly similar. In some cases, different telomeres carry the identical motifs (*e*.*g*. both telomeres of chromosome 20 in *J. angkorensis*) indicating that the original repeats could be erased and completely replaced by the spreading motif. Alternatively, novel motifs could first emerge and expand as proximal arrays, and if they provide functional solutions to the end-protection and end-replication problems, could eventually become functional telomeres. The latter idea goes in line with the hypothesis that fast-evolving subtelomeric sequences represent a driver for the divergence of both telomeric DNA sequence and corresponding DNA-binding proteins (Saint-Leandre and Levine, 2020).

It has been demonstrated that reintroduction of telomerase activity into *S. cerevisiae* type I and type II survivors restores the telomerase-dependent mode of telomere maintenance as well as the wild-type length of telomeric arrays (Teng and Zakian, 1999). Although telomerase can extend some non-canonical primers (Wang et al. 1998; Lue and Xia, 1998), it seems unlikely that it would be able to recognize diverse telomeric motifs present at the chromosomal termini of Microstromatales for addition of canonical telomeric repeats. However, the lack of tools for genetic manipulation of Microstromatales currently does not allow testing if reintroduction of telomerase and *TPP1* would restore ancestral canonical telomeres in these species.

The molecular organization of the chromosomal-end-specific telomeric arrays in Microstromatales is reminiscent of distinct telomeric repeats discovered at the ends of linear mitochondrial DNA molecules in some *Tetrahymena* species (Morin and Cech, 1988) and composite telomeres recently reported at the ends of nuclear chromosomes in nematodes of the genus *Meloidogyne* (Mota et al. 2024). Importantly, the presence of multiple types of telomeric arrays within the same cell raises questions on the mode of maintenance of individual chromosomal ends. Our searches for proteins mediating the telomere functions in Microstromatales genome sequences identified orthologs of key players participating in these processes, including those essential for ALT (**Figure 4**). Therefore, we hypothesize that noncanonical telomeres of Microstromatales are maintained via an ALT pathway similar to that operating in telomerase-deficient mutants. This idea is also supported by variable lengths of individual telomeric arrays (**Figure 2, Supplementary Figure S5**) as the heterogeneous telomere sizes represent one of the hallmarks of ALT cells (Bryan et al. 1995). It has been shown that the telomerase-deficient mutants of *M. maydis* progressively lose telomeric arrays and surviving clones emerge in senescent cultures. Such cells resemble the type I and, albeit rarely, type II survivors of *S. cerevisiae est2* mutants (Bautista-España et al. 2014; Yu et al. 2018). Since the genome sequences of *J. angkorensis* and related species lack TTAGGG-like arrays, they remind the type I survivors, which maintain the chromosomal termini by recombination mediated amplification of subtelomeric repeats such as UTASa in *M. maydis* (Sánchez-Alonso and Guzmán, 1998) or Y’ elements in *S. cerevisiae* (Louis and Haber, 1992). By analogy, the telomeric arrays of Microstromatales could arise from a similar source. Surprising variability of telomeric motifs indicates that either the ancestral chromosomes differed by subterminal repeats or these repeats underwent a rapid diversification. This goes in line with the studies showing that subtelomeres are rapidly evolving regions of the eukaryotic genomes which represent hotspots for genetic innovations (Brown et al. 2010).

## Methods

### Genome and transcriptome sequencing

Yeast strains and cultivation are described in Supplementary Information (Supplementary Methods; **Supplementary Table S1**). Yeasts from overnight cultures were harvested by centrifugation, resuspended in 100 mM EDTA pH 8.0, 2% (v/v) 2-mercaptoethanol and incubated for 30 min at room temperature. The cells were pelleted, washed and resuspended in CPES (40 mM citric acid, 120 mM Na_2_HPO_4_, 20 mM EDTA, 1.2 M sorbitol) supplemented with 5 µg/ml lysing enzymes (Sigma), 3.3 µg/ml chitinase (Sigma), 4 µl/ml Viscozyme (Sigma)) and incubated for 4.5 hours at 37 °C with occasional shaking. High-molecular weight (HMW) genomic DNA was then isolated using series of phenol and phenol: chloroform: isoamyl alcohol (25: 24: 1) extractions, precipitated with an equal volume of 96% (v/v) ethanol, washed with 70% (v/v) ethanol, and air-dried. The precipitate was dissolved in TE (10 mM Tris-HCl, 1 mM EDTA, pH 8.0) and RNA was digested by RNase A (150 μg/mL) for 30 min at 37 °C. DNA was then extracted by phenol: chloroform: isoamylalcohol (25: 24: 1), precipitated with 0.1 M NaCl and two volumes of 96% (v/v) ethanol, washed with 70% (v/v) ethanol, air-dried, dissolved in TE, and further purified by anion-exchange chromatography on a Genomic-tip 100/G (Qiagen).

For nanopore sequencing, the libraries were prepared from ~2-3 µg HMW DNA using Ligation Sequencing kits (SQK-LSK108, SQK-LSK109) and sequenced on a MinION MK-1b with a FLO-MIN106 (R9.4.1) flow cell (Oxford Nanopore Technologies) according to manufacturer’s instructions, except the fragmentation step was omitted.

Short read sequencing was performed using TruSeq DNA PCR-free (350) paired-end (2×101, 2×151-nt) libraries on Illumina platforms (HiSeq2000, NovaSeq6000) in Macrogen Europe. For RNA-Seq, yeast cultures were grown in YPD media till OD_600_ ~ 1. Total cellular RNA was extracted with hot acidic phenol (Collart and Oliviero, 1993) and purified using an RNeasy Mini kit (Qiagen). TruSeq mRNA paired-end (2×101, 2×151-nt) libraries were sequenced on a NovaSeq6000 platform in Macrogen Europe.

For telomere mapping by long read sequencing, two experiments with BAL-31 nuclease were performed. In the first experiment, two samples of *J. angkorensis* HMW DNA (~5 μg) were digested with 2 U of BAL-31 nuclease (New England Biolabs) in 50 μl of 1× BAL-31 reaction buffer at 30 °C for 15 and 30 min. In the second experiment, *J. angkorensis* HMW DNA samples (~5 μg) were digested with 1 and 3 U of BAL-31 in 50 μl of 1× BAL-31 reaction buffer at 30 °C for 30 min. In both experiments, the undigested control samples were incubated for 30 min in a 1× BAL-31 reaction buffer without the enzyme. The nuclease reactions were stopped by addition of 100 mM ethylenebis(oxyethylenenitrilo)tetraacetic acid (EGTA) pH 7.0 to a final concentration of 20 mM and the samples were incubated for 20 min at 65 °C. DNA was then purified using Agencourt AMPure XP magnetic beads (Beckman Coulter) and the tagmentation library was prepared using a Rapid Barcoding kit (SQK-RBK004) and sequenced on a MinION MK-1b with a FLO-MIN106 (R9.4.1).

All nanopore reads were basecalled by Guppy 4.4.1. Later they were basecalled by newer Dorado base caller v.0.7.2, with dna_r9.4.1_e8_sup@v3.6 model. These newer base calls were used for estimating telomere array lengths.

### Genome assemblies

For *J. angkorensis*, we assembled nanopore reads by Miniasm v.0.3-r179 (Li, 2016) and Flye v.2.8.3 (Kolmogorov et al. 2019) genome assemblers, and Illumina reads by SPAdes v.3.12.0 (Bankevich et al. 2012). The overall assembly is mostly based on the Miniasm assembly, which was polished by two iterations of Racon v1.4.3 (Vaser et al. 2017) using nanopore reads. Based on manual examination of read alignments to all three assemblies and the assemblies to each other, two contigs were connected together, some local misassemblies caused by nanopore sequencing errors were replaced by SPAdes versions and some telomeres missing in the Miniasm assembly or having wrong length were replaced by Flye versions. The resulting assembly was polished using Illumina reads by three iterations of Pilon v.1.24 (Walker et al. 2014). A single ribosomal DNA repeat was polished separately and replaced in the assembly because repetitive regions without uniquely mapping reads are difficult to polish by short reads.

The other three species (*J. pallidilutea, P. phylloscopi*, and *S. kandeliae*) were assembled by similar methods. *J. pallidilutea* is based mostly on Miniasm assembly, whereas *P. phylloscopi* and *S. kandeliae* are based mostly on Flye assemblies. SPAdes assemblies were not used in these cases. Adjustments were again made by comparing assemblies with long reads and with each other and replacing problematic pieces. Low coverage sequences were trimmed from contig ends. In *J. pallidilutea* and *S. kandeliae*, two contigs were connected manually. In *S. kandeliae*, chromosome 6 was expanded with two 20 kbp inverted repeats based on detailed read analysis. Again, Miniasm assemblies were polished by two iterations of Racon v1.4.3 using nanopore reads, the resulting assembly was polished by three iterations of Pilon using short reads and a single ribosomal DNA repeat was polished separately.

### Bioinformatic analyses of BAL-31 data

For the BAL-31 experiments, our goal was to compare sequencing coverage in the nuclease treated and control samples to determine which regions were depleted by the enzyme treatment (**Figure 1**). As read alignments are not reliable in non-unique repetitive regions, we used k-mer analysis (for k=21). First, k-mer occurrences were counted in nanopore reads using Jellyfish (Marçais and Kingsford, 2011). For each k-mer in the assembly, we tested the depletion of the k-mer in the treated sample compared to the control sample using the one-sided Fisher exact test. We selected k-mers with p-value lower than 2.5% after the Bonferroni multiple testing correction. Next, we computed the coverage of 1 kbp windows with step 100 by these significantly depleted k-mers. Windows covered more than 50% we considered as depleted regions. Nearby windows (distance less than 5 kbp) were merged to a single region. The window coverage by significant k-mers and centers of depleted regions were visualized along chromosomes using the ggplot2 library in R (Wickham, 2011).

### Telomeric motifs

Telomeric and subtelomeric motifs were found by Tantan (Firth, 2011) with maximum motif length set to 250 bp in the terminal 50 kbp at each chromosomal contig end. The found motifs and their occurrences were then examined manually to exclude other tandem repeats located further from telomeres and to avoid redundant overlapping motifs. All motifs are listed in the 5’ to 3’ direction leading out of the chromosome (that is, motifs from left chromosomal ends are reversed compared to their appearance in the assembly).

To estimate the lengths of telomeric arrays, we selected nanopore reads that span the unique sequence region nearest to the chromosome end and counted the number of bases covered by telomeric and subtelomeric motifs. To lessen the impact of the reads that end before the chromosome end, we selected only reads longer than a threshold selected based on read length and coverage in a particular species (30 kbp for *J. angkorensis* and *J. pallidilutea*, 20 kbp for *P. phylloscopi*, 25 kbp for *S. kandeliae*). To search for telomeric motifs in reads, we created a hidden Markov model (HMM) with a single background state representing non-repetitive sequence and a simple profile HMM (Durbin et al. 1998) for each distal and proximal motif found at a particular chromosomal end (see details in Supplementary Methods and **Supplementary Figure S8**). HMMs are well suited for modeling tandem repeats (Firth, 2011, Nánási et al. 2014, Olson and Wheeler, 2024). The most probable sequence of states for a given read segment was estimated using the standard Viterbi algorithm (Durbin et al. 1998). A typical read consists of a subtelomeric segment emitted by the background state, followed by one or several telomeric segments emitted by the profile HMMs and a short background segment located at the very end corresponding to adapters or low-quality sequence. Spans of lower quality sequence emitted by the background state can interrupt telomeric repeats. We report the total number of bases emitted by the profile HMM states as the length of the telomeric array in a given read. We show the violin plot of these lengths, and we use the third quartile as our estimate of the length for a given chromosomal end. We use the third quartile instead of the median, as some of the reads do not extend to the full length of the chromosome.

To assess similarity of motif sequences, we enumerated all 7-mers in each motif (including 7-mers spanning the boundary between successive motif copies) and compared 7-mer multisets by the containment distance defined as 1-N_i_/N_s_ where N_i_ is the size of the intersection of the 7-mer multisets for the two motifs and N_s_ is the size of the smaller multiset. The distance is always between 0 and 1, with value 0 if the multisets are identical or one is a subset of the other and value 1 if they have no k-mer in common. We applied the single-linkage agglomerative clustering, where we merge clusters until a given distance threshold, with distance between clusters being the distance between nearest cluster representatives. We show clusters as distance thresholds 0.1, 0.7, and 0.8.

## Data availability

The genome assemblies, Illumina, and nanopore reads have been deposited in the European Nucleotide Archive (ENA) and GenBank databases. The accession numbers are shown in **Supplementary Table S1**. The genomes can be interactively visualized in the genome browser at http://genome.compbio.fmph.uniba.sk/.

## Code availability

Software developed for the analysis in this article is available at https://github.com/fmfi-compbio/jamang-code. Existing software tools are listed in the Methods section.

## Author contributions

Conceptualization, methodology, supervision, and funding acquisition: B.B., L.T., T.V., and J.N.; Experimental work: V.H., H.L., F.B., F.C., T.P., E.H., M.F.J., and J.N.; Bioinformatic analyses: B.B., A.G., D.B., T.V., and J.N.; Writing the manuscript draft: B.B. and J.N.; Review and editing: B.B., V.H., F.B., F.C., T.P., E.H., M.F.J., M.N., L.T., M.S., T.V., and J.N. All authors approved the final version.

## Materials & Correspondence

to Broňa Brejová and Jozef Nosek.

## Acknowledgments

This research was supported by grants from the Slovak Research and Development Agency (18-0239 and 22-0144 (to J.N.), 23-0056 (to L.T.)), the Scientific Grant Agency of the Ministry of Education, Science and Sport of the Slovak Republic (1/0538/22 (to T.V.), 1/0234/23 (to J.N.), 1/0031/24 (to L.T.), and 1/0140/25 (to B.B.)). Additional support was provided by the Advancing University Capacity and Competence in Research, Development and Innovation (ACCORD) project and the European Union NextGenerationEU through the Recovery and Resilience Plan for Slovakia under the project No. 09I03-03-V06-00079 (to J.N.).

