## Supplementary Information for "Noncanonical chromosomal-end-specific telomeric arrays in naturally telomerase-negative yeasts"

### Supplementary Methods

**Yeast strains and cultivation.** The type strain of *Jaminaea angkorensis* C5b (CBS 10918) was from the yeast collection of M. Sipiczki (Sipiczki and Kajdacsi, 2009). *Jaminaea pallidilutea* CBS 14684 (Nasr et al. 2017)*, Parajaminaea (Jaminaea) phylloscopi* CBS 14087 (Francesca et al. 2016), and *Sympodiomycopsis kandeliae* CBS 11676 (Wei et al. 2011) were purchased from the Westerdijk Fungal Biodiversity Institute (Utrecht, The Netherlands; **Supplementary Table S1**). Yeast cultures were grown in liquid YPD media (1% (w/v) yeast extract, 2% (w/v) peptone, 2% (w/v) glucose) at 28 $^{o}$C with constant aeration. YM medium (0.3% (w/v) yeast extract, 0.3% (w/v) malt extract, 0.5% (w/v) peptone, 1% (w/v) glucose, 2% (w/v) agar) was used for cultivation on plates (**Supplementary Figure S9**).

**Telomere restriction fragments (TRF) analysis.** Total cellular DNA of *J. angkorensis* was digested with endonucleases *Eco*RV, *Hin*dIII (New England Biolabs), and *Xho*I (Fermentas), according to the manufacturers' instructions (~3 U/µg of DNA). The reactions were carried out overnight at 37 °C. Digested DNA samples (~1 µg) were electrophoretically separated in a 0.7% (w/v) agarose gel in 0.5×TBE (45 mM Tris-borate, 1 mM ethylenediaminetetraacetic acid (EDTA), pH 8.0) for about 4 hours at 5 V/cm. The DNA in gels was depurinated (5 min in 0.25 N HCl), denatured (40 min in 1.5 M NaCl, 0.5 M NaOH), and neutralized (30 min in 0.5 M Tris-HCl, 1.5 M NaCl, 1 mM EDTA, pH 7.5). The depurination step was omitted for analysis of short fragments. The gels were then soaked for 30 min in 20×SSC (3 M NaCl, 0.3 M sodium citrate, pH 7.0) and DNA was transferred overnight onto an Immobilon-Ny+ membrane (Millipore) using a TurboBlotter system (Schleicher & Schuell). After the transfer, the membrane was rinsed with 2×SSC, air-dried and baked for 90 min at 80 °C. The membrane strips were pre-hybridized in 5×SSC, 5×Denhardt's solution, 0.5% (w/v) sodium dodecyl sulfate (SDS) for 90 min at 65 °C and then incubated overnight in a hybridizer (Techne) at 60 °C in the same buffer supplemented with 1.2 nM oligonucleotide probe (**Supplementary Table S4**) radiolabeled using a T4 polynucleotide kinase (Thermo Fisher Scientific) and [$\gamma^{32}$P]-ATP (3000 Ci/mmol, Hartmann Analytic), and purified by Illustra MicroSpin G-25 columns (GE Healthcare). The membrane strips were washed once with 2×SSC, 0.1% (w/v) SDS for 5 min at room temperature and with 0.5×SSC, 0.1% SDS two times for 15 min at 55 °C. The membranes were exposed to a phosphor screen and the signal was detected by Personal Molecular Imager FX (Bio-Rad). The contrast of images was adjusted using Quantity-One v. 4.6 software (Bio-Rad).

**Sensitivity of telomeric DNA to BAL-31 nuclease.** Total cellular DNA of *J. angkorensis* (~4.5 µg) was digested with 0, 3, 6, 12, and 24 U of BAL-31 nuclease (New England Biolabs) for 30 min at 30 °C in a final volume 120 µl of 1× BAL-31 reaction buffer. The reaction was stopped by addition of ethylenebis(oxyethylenenitrilo)tetraacetic acid (EGTA) to 30 mM and heated for 10 min at 65 °C. DNA was then purified using AMPure XP beads (Beckman Coulter) and digested with restriction endonuclease *Eco*RV (New England Biolabs). Agarose gel electrophoresis and Southern blot hybridization were performed essentially as described above.

**Analysis of single-stranded telomeric DNA.** Total cellular DNA samples (75 ng each) of *J. angkorensis* were digested in 50 µl reactions using 26.7 U of exonuclease I (ExoI, New England Biolabs), 12 U of $RecJ_f$ (New England Biolabs), and 13.3 U of mung bean nuclease (MBN, New England Biolabs) at 37 °C for 3 hours (ExoI, $RecJ_f$) or at 30 °C for 45 min (MBN). The volumes of native nuclease treated as well as undigested control samples were adjusted to 100 µl with 1×SSC. For denatured samples, DNA in 50 µl of 0.2 M NaOH was incubated at 65 °C for 15 min and the final volume adjusted to 100 µl with 1×SSC. Dot blotting was performed using Immobilon Ny+ membrane (Millipore) and DHM-48 plastic manifold (Scie-Plas) under 80 mbar vacuum. The crosslinking was performed in a CX-2000 UV crosslinker (UVP) for 18 s at 5000 µJ/cm2. The membrane strips were pre-hybridized in 5×SSC, 5×Denhardt solution, 0.5% (w/v) SDS at 47.5 °C for 1 h, and then hybridized overnight with labeled oligonucleotide probes (**Supplementary Table S4**) in the same buffer at 47.5 °C. The membranes were washed for 15 min with 5×SSC; 0.1% (w/v) SDS at 25 °C, followed by two more washes at 42 °C. Following the exposure to a phosphor storage screen, image acquisition was performed with Amersham Typhoon scanner.

**Electrophoretic karyotyping.** Chromosomal DNA samples were prepared from yeast cells embedded in agarose plugs as follows. About $8×10^8$ cells from an overnight culture grown in YPD medium (1% (w/v) yeast extract, 2% (w/v) peptone, 2% (w/v) glucose) were washed successively with water, 50 mM EDTA pH 8.0 and CPES buffer (1.2 M sorbitol, 40 mM citric acid, 120 mM $Na_2HPO_4$, 20 mM EDTA pH 8.0). The cells were then resuspended in 1 mL of CPES buffer supplemented with 5 mM dithiothreitol, 15 µg chitinase (Sigma), 20 µl viscozyme (Sigma), and 24 µg lysing enzyme (Sigma), and incubated for 60 min at 37 ºC. Spheroplasts were pelleted by centrifugation (1700×*g* for 1 min), resuspended in 1 ml of 1% (w/v) low melting point agarose (UltraPure, Life Technologies) prepared in 50 mM EDTA pH 8.5 and incubated in 10 mM Tris-HCl pH 7.5, 0.45 M EDTA pH 8.5 for 30 min at 37 ºC. The buffer was then replaced by 0.45 M EDTA pH 8.5, 1% (w/v) N-lauroyl sarcosine, 1 mg/mL proteinase K (Applichem) and incubated for 48 hours at 50 ºC and the plugs were washed with 0.45 M EDTA pH 8.5, 1% (w/v) N-lauroyl sarcosine. Chromosomal DNAs were separated by PFGE in a Pulsaphor unit (LKB) with a contour-clamped homogeneous electric field configuration (CHEF) as specified in legend to **Supplementary Figure S1.**

**Genome annotations.** To annotate genomes, we assembled RNA-seq reads into transcripts by Trinity v2.15.1 (Grabherr et al. 2011). Before assembly, reads were trimmed by Trimmomatic v.0.36 (Bolger et al. 2014). The resulting transcripts were aligned to the corresponding assembly by Blat v.36x7 (Kent 2002) with option -maxIntron=5000. Using pslCDnaFilter from UCSC browser utilities (Kuhn et al 2013), we filtered out alignments with identity less than 95% or covering less than 75% of the transcript and kept only the best alignment for each transcript. Protein coding genes were predicted by Augustus v.3.2.3 (Stanke et al. 2006) with transcript alignments used as hints supporting exon positions. For *J. angkorensis*, we started from parameters provided with Augustus for *M. maydis*. After predicting genes, we selected genes supported by RNA-seq transcripts and trained new parameters adjusted for this genome on these high-confidence genes. Namely, we aligned the predicted genes to the assembled transcripts by blat and kept only alignments covering 99% of the predicted genes with at least 99% sequence identity and no gaps. Genes having such alignments were used for parameter training. The trained parameters were then again used in Augustus for final prediction. For the other three species, we started from parameters trained on *J. angkorensis* and retrained it on the respective genome using the same method. For *S. kandeliae*, two rounds of training were performed, because the initial gene prediction yielded only a small number of supported genes for training (2694 supported genes after the initial gene predictions, 5141 after the the first iteration). This was likely due to the large difference in genome GC content between these species (49.8% for *S. kandeliae* and 60.3% for *J. angkorensis*). The resulting gene predictions and supporting evidence from RNA-seq alignments were visualized in the UCSC genome browser (Kuhn et al. 2013). Selected genes were inspected manually, and predictions were corrected based on other evidence (RNA-seq, protein homology).

The reference genome and predicted proteome sequences of *J. angkorensis*, *J. pallidilutea*, *P. phylloscopi*, and *S. kandeliae* generated in this study and the seven Exobasidiomycete species (*Acaromyces ingoldii* (GCF_003144295.1), *Ceraceosorus guamensis* (GCF_003144195.1), *Jaminaea rosea* (GCF_003144245.1), *Meira miltonrushii* (GCF_003144205.1), *Pseudomicrostroma glucosiphilum* (GCF_003144135.1), *Tilletiaria anomala* (GCF_000711695.1), and *Tilletiopsis washingtonensis* (GCF_003144115.1)) downloaded from the GenBank database (https://www.ncbi.nlm.nih.gov/datasets/genome/; Kijpornyongpan et al. 2018; Toome et al. 2014) were used in sequence analyses. Amino acid sequences of *M. maydis* proteins downloaded from the Uniprot database (https://www.uniprot.org/; UniProt Consortium, 2023) were used as queries in BlastP and tBlastN searches (2.15.0+; Altschul et al. 1997) of the predicted proteomes and reference genome assemblies, respectively, with an expect value (e-value) threshold 1e-1. Identified proteins were next used as queries in reciprocal Blast searches against the *M. maydis* proteome. In addition, the searches of HMM domain profiles were performed using hmmsearch (HMMER 3.4; http://hmmer.org/; Eddy, 2011). Note that the homologs of Ten1 (*J. angkorensis, J. pallidilutea, P. phylloscopi*) and Trf1/Tay1 (*P. phylloscopi*) are weakly conserved and were not detected in the Blast searches using the *M. maydis* queries. Ten1 in Microstromatales was identified by the presence of the Ten1_2 domain as well as by the BlastP searches with the *S. kandeliae* Ten1 query. Trf1/Tay1 in *P. phylloscopi* was searched using the *J. angkorensis* Trf1/Tay1 query and confirmed by the presence of two SANT/Myb domains.

To visualize homology between *J. angkorensis* and the other three genomes (**Supplementary Figure S6**), genome assemblies were aligned using Last aligner v.1542 (Frith et al. 2010) with E-value threshold E1e-10. The alignments were post-processed by last-split utility, which ensures that each portion of *J. angkorensis* is covered by at most one alignment of the other genome. The alignments were then visualized by a custom script.

To explore tandem repeat arrays in whole genomes (**Supplementary Figure S7**), Tantan v.22 (Firth, 2011) was run on the whole genome sequences of the four studied species as well as the genome of *M. maydis* (NCBI RefSeq assembly GCF_000328475.2), searching for repeats of length at most 250. Only motifs of length at least 30 with at least 2 full repeats were kept.

**Hidden Markov model for finding telomeric repeats.** Each chromosomal end in each studied species is characterized by one or several motifs (proximal and distal) (**Supplementary Table S3**). From these motifs we build an HMM consisting of a single background state and a profile HMM for each motif (**Supplementary Figure S8**). The profile HMMs are circularized with the first state following the last state to model tandem repeat nature of the repeats. From the background state, the model can enter any of the profile HMM states with equal probability; the overall probability of leaving the background state and entering one of the motifs was set to 0.001. Similarly, the model can enter the background state from any motif state with probability 0.0001. The profile hidden Markov models for individual motifs have fixed probabilities of mismatch (0.01), insertion opening (0.05) and deletion opening (0.05) at each position. The probability of extending insertion and deletion was set to 0.5 and the maximum deletion length was capped at 7 bp. The mismatches, insertions and deletions may represent sequencing errors in nanopore reads, but also any biological variation in these repeats.

**Phylogenetic analysis.** The phylogeny (**Figure 4**) was inferred from the genome sequences (see above). Single-copy orthologs were identified in the genome sequences by BUSCO v. 5.1.2 in the genome mode with basidiomycota_odb10 lineage (Manni et al. 2021) and the supermatrix was calculated from 1,764 orthologs present in at least 50% of the genomes using the BUSCO_phylogenomics pipeline (options: –psc 50 –supermatrix; McGowan, 2023). The tree was built using FastTree 2.1.11 (Price et al. 2010) and the ggtree package in R 4.3.1 (Yu et al. 2017).

**Supplementary Table S1A.** Yeast strains.

| Species | Strain | Strain status | Source | Origin |
|---|---|---|---|---|
| *Jaminaea angkorensis* | C5b (CBS 10918) | Type | M. Sipiczki | from fallen leaves (Angkor, Cambodia) |
| *Jaminaea pallidilutea* | CBS 14684 | Holotype | Westerdijk Fungal Biodiversity Institute, The Netherlands | from leaves of *Avicennia marina* (Qeshm Island, Iran) |
| *Parajaminaea (Jaminaea) phylloscopi* | CBS 14087 | Holotype | Westerdijk Fungal Biodiversity Institute, The Netherlands | from a trans-saharan migratory bird (Ustica Island, Italy) |
| *Sympodiomycopsis kandeliae* | CBS 11676 | Type | Westerdijk Fungal Biodiversity Institute, The Netherlands | from a flower of *Kandelia candel* (Hsinchu, Sinfong, mangrove forest, Taiwan) |

**Supplementary Table S1B.** Nanopore and Illumina reads.

| Species | ONT reads (DNA) | | | | | Illumina reads (DNA) | | | | | Illumina reads (RNA) | | | |
|---|---|---|---|---|---|---|---|---|---|---|---|---|---|---|
| | Number | [Gbp] | N50 [kbp] | Coverage | ENA acc. no. | Number | [Gbp] | paired-end | Coverage | ENA acc. no. | Number | [Gbp] | paired-end | ENA acc. no. |
| *Jaminaea angkorensis* | 557,199 | 6.52 | 20.0 | 314x | ERR15532040 | 65 392 686 | 6.60 | 2x101 | 318x | ERR15532041 | 45 098 496 | 4.55 | 2x101 | ERR15532042 |
| *Jaminaea pallidilutea* | 536,248 | 2.66 | 13.0 | 154x | ERR15532034 | 8 572 620 | 1.29 | 2x151 | 74x | ERR15532035 | 38 233 536 | 5.77 | 2x151 | ERR15532036 |
| *Parajaminaea (Jaminaea) phylloscopi* | 769,184 | 2.98 | 10.2 | 162x | ERR15532037 | 49 253 342 | 7.44 | 2x151 | 406x | ERR15532038 | 41 243 012 | 6.23 | 2x151 | ERR15532039 |
| *Sympodiomycopsis kandeliae* | 663,324.0 | 2.96 | 14.0 | 153x | ERR15532030 | 48 083 632 | 7.26 | 2x151 | 376x | ERR15532032 | 42 067 610 | 6.35 | 2x151 | ERR15532033 |
| | 335,026.0 | 1.14 | 9.0 | 59x | ERR15532031 | | | | | | | | | |

**Supplementary Table S1C.** Nanopore reads from BAL-31 experiments.

| ***J. angkorensis*** | Sample | ONT reads (DNA) | | | | |
|---|---|---|---|---|---|---|
| | | Number | [Gbp] | N50 [kbp] | Coverage | ENA acc. no. |
| Experiment #1 | control 1 (w/o BAL-31), 30 min | 599,861 | 1.72 | 5.0 | 83x | ERR15532043 |
| | BAL-31 (0.2 U/μg DNA), 15 min | 460,121 | 1.27 | 4.7 | 61x | ERR15532044 |
| | BAL-31 (0.2 U/μg DNA), 30 min | 323,461 | 0.77 | 3.9 | 37x | ERR15532045 |
| Experiment #2 | control 2 (w/o BAL-31), 30 min | 427,319 | 1.92 | 9.0 | 92x | ERR15532046 |
| | BAL-31 (0.2 U/μg DNA), 30 min | 66,53 | 0.29 | 8.5 | 14x | ERR15532047 |
| | BAL-31 (0.6 U/μg DNA), 30 min | 326,317 | 1.52 | 8.4 | 73x | ERR15532048 |

**Supplementary Table S1D.** Characteristics of the genome assemblies.

| Species | Strain | Chromosomal contigs | Nuclear genome size [Mbp] | Nuclear genome %G+C | Predicted proteins | Bioproject [acc.no.] | mtDNA [bp] | mtDNA [%G+C] | mtDNA [acc. no.] | BUSCO v. 5.1.2 #genome mode | | | | BUSCO v. 5.1.2 #proteins mode | | | |
|---|---|---|---|---|---|---|---|---|---|---|---|---|---|---|---|---|---|
| | | | | | | | | | | Complete and single-copy | Complete and duplicated | Fragmented | Missing | Complete and single-copy | Complete and duplicated | Fragmented | Missing |
| *Jaminaea angkorensis* | C5b (CBS 10918) | 20 | 20.75 | 60.3 | 8 263 | PRJEB35162 | 29 999 | 32,2 | KC628747; Hegedusova *et al*. 2014 | 90.0 | 0.2 | 1.9 | 7.9 | 91.5 | 0.2 | 1.0 | 7.3 |
| *Jaminaea pallidilutea* | CBS 14684 | 16 | 17.27 | 55.4 | 6 317 | PRJEB77303 | 43 965 | 25,3 | PP995848 | 90.6 | 0.1 | 1.7 | 7.6 | 91.6 | 0.1 | 0.7 | 7.6 |
| *Parajaminaea (Jaminaea) phylloscopi* | CBS 14087 | 3 | 18.32 | 61.0 | 6 583 | PRJEB77304 | 49 037 | 28,6 | PP995849 | 91.5 | 0.1 | 1.4 | 7.0 | 92.6 | 0.1 | 0.9 | 6.4 |
| *Sympodiomycopsis kandeliae* | CBS 11676 | 6 | 19.29 | 49.8 | 7 053 | PRJEB77302 | 39 719 | 27,6 | PP995850 | 91.4 | 0.3 | 1.6 | 6.7 | 92.3 | 0.4 | 0.9 | 6.4 |

BUSCO genome mode: Lineage dataset: basidiomycota_odb10 (2024-01-08, number of genomes: 133, number of BUSCOs: 1764)
Gene predictor: metaeuk

BUSCO proteins mode: Lineage dataset: basidiomycota_odb10 (2024-01-08, number of genomes: 133, number of BUSCOs: 1764)

**Supplementary Table S2A.** Characteristics of telomeric arrays in *J. angkorensis*.

| Chromosome | | | | Left Telomere | | | Right Telomere | | |
|---|---|---|---|---|---|---|---|---|---|
| **No.** | **Size [bp]** | **G+C%** | **Median coverage** | **Proximal repeat motif [bp]** | **Distal repeat motif [bp]** | **Array size estimate [kbp]** | **Proximal repeat motif [bp]** | **Distal repeat motif [bp]** | **Array size estimate [kbp]** |
| **01** | 2 873 230 | 60.4 | 303 | | 72 | 2.0 | | 73 | 8.6 |
| **02** | 2 288 400 | 60.4 | 303 | | 167 | 8.6 | 80 | 52[a] | 5.6 |
| **03** | 1 591 733 | 60.4 | 304 | | 73 | 18.0 | | 72 | 13.5 |
| **04** | 1 373 946 | 59.8 | 301 | | 72 | 4.8 | 73 | 178[b] | 12.8 |
| **05** | 1 282 165 | 60.4 | 303 | | 73 | 18.1 | | 59 | 20.4 |
| **06** | 1 121 287 | 59.9 | 298 | | 73 | 7.3 | | 73 | 3.0 |
| **07** | 1 106 543 | 60.6 | 299 | | 63 | 15.2 | | 73 | 12.2 |
| **08** | 907 346 | 60.6 | 300 | | 62 | 8.4 | 126 | 52[a] | 19.7 |
| **09** | 883 201 | 60.7 | 293 | | 100 | 6.0 | | 156 | 4.4 |
| **10** | 847 713 | 60.3 | 294 | | 72 | 17.1 | 155 | 137 | 18.1 |
| **11** | 831 147 | 60.4 | 296 | | 152 | 3.8 | | 73 | 9.5 |
| **12** | 822 261 | 60.4 | 291 | | 153 | 4.3 | | 83 | 16.4 |
| **13** | 799 268 | 60.2 | 300 | | 73 | 4.4 | 72 | 52[a] | 10.6 |
| **14** | 770 973 | 60.6 | 298 | | 63 | 5.9 | | 72 | 3.7 |
| **15** | 759 781 | 60.6 | 300 | | 73 | 1.8 | | 72 | 1.6 |
| **16** | 620 785 | 60.3 | 295 | | 72 | 17.1 | | 59 | 21.6 |
| **17** | 596 954 | 59.9 | 301 | | 151 | 5.6 | | 72 | 2.9 |
| **18** | 563 986 | 60.6 | 289 | | 83 | 5.7 | 63 | 178[b] | 20.5 |
| **19** | 545 337 | 60.4 | 298 | | 74 | 19.0 | | 164 | 6.2 |
| **20** | 135 699 | 59.2 | 291 | | 130[c] | 4.2 | | 130[c] | 7.0 |

a, b or c - these arrays contain identical motifs

**Supplementary Table S2B.** Characteristics of telomeric arrays in *J. pallidilutea*.

| Chromosome | | | | Left Telomere | | | Right Telomere | | |
|---|---|---|---|---|---|---|---|---|---|
| **No.** | **Size [bp]** | **G+C%** | **Median coverage** | **Proximal repeat motif [bp]** | **Distal repeat motif [bp]** | **Array size estimate [kbp]** | **Proximal repeat motif [bp]** | **Distal repeat motif [bp]** | **Array size estimate [kbp]** |
| **01** | 3 600 319 | 55.6 | 70 | | 96 | 2.8 | | 81[f] | 5.9 |
| **02** | 2 240 588 | 55.5 | 69 | 56[c], 57 | 100[h] | 4.8 | | 63 | 3.9 |
| **03** | 1 202 730 | 55.1 | 67 | | 41[a] | 7.9 | 58[e] | 100[h] | 5.6 |
| **04** | 1 092 581 | 55.7 | 67 | | 96[g] | 3.6 | 60 | 81[f] | 4.9 |
| **05** | 1 084 863 | 55.4 | 67 | | 55 | 3.8 | 56 | 100[h] | 7.5 |
| **06** | 1 083 760 | 55.5 | 67 | | 42[b] | 3.2 | 57[d] | 100[h] | 5.8 |
| **07** | 913 777 | 55.1 | 66 | | 96 | 5.0 | 43 | 100[h] | 5.7 |
| **08** | 858 994 | 55.5 | 67 | | 62 | 5.0 | | 96[g] | 3.7 |
| **09** | 856 032 | 55.3 | 66 | 58[e] | 100[h] | 3.4 | | 84 | 6.4 |
| **10** | 806 894 | 55.5 | 67 | 57[d] | 96 | 3.8 | 56 | 100[h] | 5.4 |
| **11** | 709 386 | 55.3 | 129 | | 42[b] | 2.2 | | 83 | 7.8 |
| **12** | 707 902 | 55.4 | 65 | | 75 | 2.4 | 58[e] | 99 | 6.7 |
| **13** | 680 837 | 55.3 | 67 | | 81 | 1.5 | 56[c] | 96[g] | 4.1 |
| **14** | 658 102 | 55.5 | 66 | | 41[a] | 6.9 | 56, 96, 58[e], 99 | 100[h] | 5.4 |
| **15** | 471 378 | 55.5 | 65 | 101 | 100[h] | 4.5 | | 81 | 5.1 |
| **16** | 257 273 | 54.1 | 64 | | 46 | 3.2 | | 41[a] | 4.7 |

a, b, c, d, e, f, g or h - these arrays contain identical motifs

**Supplementary Table S2C.** Characteristics of telomeric arrays in *P. phylloscopi*.

| Chromosome | | | | Left telomere | | | Right telomere | | |
|---|---|---|---|---|---|---|---|---|---|
| **No.** | **Size [bp]** | **G+C%** | **Median coverage** | **Proximal repeat motif [bp]** | **Distal repeat motif [bp]** | **Array size estimate [kbp]** | **Proximal repeat motif [bp]** | **Distal repeat motif [bp]** | **Array size estimate [kbp]** |
| **01** | 9 859 025 | 60.8 | 392 | | 121 | 8.6 | | 132 | 9.8 |
| **02** | 4 562 156 | 61.3 | 384 | | 101 | 6.4 | 101 | 130[a] | 18.2 |
| **03** | 3 850 713 | 61.1 | 375 | 81 | 130[a] | 7.1 | | 101 | 19.2 |

a - these arrays contain identical motifs

**Supplementary Table S2D.** Characteristics of telomeric arrays in *S. kandeliae*.

| Chromosome | | | | Left Telomere | | | Right Telomere | | |
|---|---|---|---|---|---|---|---|---|---|
| **No.** | **Size [bp]** | **G+C%** | **Median coverage** | **Proximal repeat motif [bp]** | **Distal repeat motif [bp]** | **Array size estimate [kbp]** | **Proximal repeat motif [bp]** | **Distal repeat motif [bp]** | **Array size estimate [kbp]** |
| **01** | 5 778 576 | 49.8 | 317 | 69 | 62[a] | 30.5 | | 70[b] | 21.7 |
| **02** | 4 498 260 | 49.7 | 311 | 111 | 111 | 20.7 | 69, 62 | 62[a] | 22.6 |
| **03** | 3 765 011 | 49.8 | 319 | | 70[b] | 18.3 | | 62[a] | 17.4 |
| **04** | 3 138 396 | 49.9 | 315 | | 62 | 20.3 | | 61 | 24.9 |
| **05** | 1 870 769 | 49.9 | 306 | | 122 | 23.4 | | 70[b] | 23.5 |
| **06** | 198 920 | 51.8 | 276 | | 55[c] | 18.3 | | 55[c] | 20.9 |

a, b or c - these arrays contain identical motifs

**Supplementary Table S3A.** Telomeric sequences of *J. angkorensis*

| Chromosome / position | Size [bp] | G+C% | # reads | median [bp] | Q3 [bp] | Note | Motif [5'->3'] |
|---|---|---|---|---|---|---|---|
| **Proximal repeats** | | | | | | | |
| 2 / Right | 80 | 57.5 | 99 | 2302.0 | 2354.5 | | GGGGGCAGGGAAGGTGTGGTTTGGTGTGGGGTTTGTGATTGGGAGCGTAGCGGTGGTGTGATTGTTGGAGTGGTCTGTCA |
| 4 / Right | 73 | 58.9 | 114 | 3229.0 | 3371.0 | | GGGTTGCAGCGTATCTGGGCCTTTTGGTGCACAGCTTGGCGATCGAGGCCCAGGAAAGGGTGCTTGTACGGTT |
| 8 / Right | 126 | 60.3 | 57 | 8683.0 | 9109.0 | | GGGGTGGGTGTGGCTCTTTGGCAGCGTAGCTTGTAGTGCGAGGGCCAGGAAAGGGTGTGGTGTGGTGTTGATTGGGAGCGTAGCTGTCCT<br>CTGTCTGGCGTGGATCGTAGTGCGAGGGCCAGGAAA |
| 10 / Right | 155 | 54.2 | 118 | 2073.5 | 2122.8 | | GGGGCTTTTAGTGTGTGGTAGTGTCCAGCATTGGTTCTCTTGCGTAATAGGAAAGCCAGTACGGCATGATTGGCCTGTGGTACGGTGTATAG<br>GTGGATGAGGTGGTGTGTTTGGCAGTGTCTCTGGCTGGCGTGGTTGCCTGGTTGGCGAGCTAC |
| 13 / Right | 72 | 62.5 | 78 | 6564.5 | 6794.8 | | GGGGCGGGAAAGGGGTGAGTGTGGCTTTGTGGGAGCGTAGGTGGCTTCCGTCTGGCGTAGCTTGTAGTGCGA |
| 18 / Right | 63 | 61.9 | 114 | 793.0 | 811.5 | | GGGGGATCAGGAAAGGCGTGGGTTGTGCACTTGCTGTGCGGTTGCTCTGCGCTTGTGGCGTAC |
| **Distal repeats** | | | | | | | |
| 1 / Right end | 73 | 63.0 | 103 | 8341.0 | 8600.5 | | GGGGGCTGCGTGGCCTCGTGTGCTGCGTGTGGCTACGGTAGCCGATCTCATCTGACTGAGGAAAGAGGGCGTA |
| 1 / Left end | 72 | 59.7 | 91 | 1838.0 | 1960.0 | | GGGGCTGGCTTTTGTTGTGAGCGGCGTGGGTGCGTGGCACTGGTCCGTCGAGTTCTCATTTGGGTCTGGTAA |
| 2 / Right end | 52 | 61.5 | 99 | 3051.0 | 3248.5 | (a) | GGGGTGGGTGTGGCTCTTTGGCAGCGTAGCTTGTAGTGCGAGGGCCAGGAAA |
| 2 / Left end | 167 | 47.9 | 104 | 8449.0 | 8603.0 | | GGGAAAGGGCTGAGTGGTGTGTGGTTGGTGATGGACCTTTTAGTCTATTGCGTGCTAGGAAAGCAGAATGCACTTGATCTTGATGGCTAGA<br>GTTGTAGGGCACAGTGAGAGCTTTGCAATGATACCATTTAGGTCCTGATTGGGTGAATACTGGCGGAGGTATTGGT |
| 3 / Right end | 72 | 52.8 | 96 | 10579.5 | 13529.3 | | GGTGGATCAGGAAAGATATGGTGTGTGGTTTTGAGTGGCGTACGACCTTGCTGTGCGCCGTTGCTTTGCTCT |
| 3 / Left end | 73 | 54.8 | 96 | 17204.0 | 17956.3 | | GGGCTGCTGTAGCCGTAACTGCTCTGTCAAGGCACAGGAAAGGGTCATGGTATGGTTTGCTTGTGGCGTACCT |
| 4 / Right end | 178 | 60.1 | 114 | 8922.5 | 9434.3 | (b) | GGGGGATCAGTGCTAGCGTGGGTGCCCTCTAGGAATTAGGAAAGAGTGAGTGACGGATTGCGAGCTAGGGAAGGTATCTGGGCCTTTTGG<br>TGCACAGCTTGGCGATCGAACCACCGGCGTAGCGTAGCCGGCGGTGGTGACTACCGGCGGATAGGCAAGTGGTGCGCTTGTGGCGTAC |
| 4 / Left end | 72 | 58.3 | 125 | 4657.0 | 4839.0 | | GGCTTGGTGTGGCGTAGCCCGTGTGCAGTCTCCGTAGCTGTCTCAGCAAGGCCCAGGAAAGTGTGAGTGTAT |
| 5 / Right end | 59 | 57.6 | 97 | 19076.0 | 20441.0 | | GGGGTAAGGGTTGAGCCTGTTATTTCTGGCGGCGAGGTGGGAAAGCCCTTTGGTTCCTC |
| 5 / Left end | 73 | 57.5 | 75 | 17469.0 | 18131.5 | | GGGGCTTTTGAGCAGCGTAGCCGTTGATTTGGCTGGGTGTTTGTTCTGGCGTGGCCCAGGAAAGGGCTGTTGT |
| 6 / Right end | 73 | 63.0 | 105 | 2946.0 | 3041.0 | | GGGCCAGGAAAGCCAGCGTACGGCAACCGTAGGGCTGCCTGTGGCTCCGTAGCTGTGGTTCTGTGTCTGACCA |
| 6 / Left end | 73 | 60.3 | 97 | 7064.0 | 7282.0 | | GGGCGTAGGAAAGGGCTTGTAAGGCGCTGCGGACCAGCGCACTTGTGGTTTGCTGAGCATGGCTGACTTGCCA |
| 7 / Right end | 73 | 58.9 | 98 | 11901.5 | 12212.8 | | GGGGGGAAAGCCTGGCTGTATGGTTTCCTCCTAGCGTAGCTGGCTGCTTGGTGGCGTCATTCACCTACTGCGT |
| 7 / Left end | 63 | 55.6 | 103 | 14607.0 | 15195.0 | | GGGCCCCAGAACCGGCTTTGGTTGGTCGATCTACTGCGAGATAGGAAAGTGTGGTGTGCTTGT |
| 8 / Right end | 52 | 61.5 | 57 | 10031.0 | 10560.0 | (a) | GGGGTGGGTGTGGCTCTTTGGCAGCGTAGCTTGTAGTGCGAGGGCCAGGAAA |
| 8 / Left end | 62 | 62.9 | 51 | 8175.0 | 8389.0 | | GGGGGCTGTCTCCTTCTACGGCCTCACATGGGTCAGGAAAGGGCGTGCTGTGGTGCGTAGCT |
| 9 / Right end | 156 | 58.3 | 86 | 4294.5 | 4405.3 | | TTGTCTTGTCCGGCGTATTCCGCCGTAGCCGATTGGCTACTGTTTGCTGGTGCTTGACTTCTGGGCGGGAAAGTGGCTGCTTTTGGCCTGCTT<br>CTGGTGTGAATGGGTGTAGCCGTGGGTGGGCTCTGAGTGGTGGGAGTTGTGTGTGGCGTGGCT |
| 9 / Left end | 100 | 57.0 | 95 | 5840.0 | 5998.5 | | GGGGGACCTTGGTCTTGACTGTGCTTAGCAGCCGTCTTGTGGCGAGGGTTAGGAACGAGCAGCTTAGTGACTTGCTTGGCGAGTGCTGACG<br>CCAAAGAGT |
| 10 / Right end | 137 | 56.2 | 118 | 15317.5 | 15958.5 | | GGGGATCGAGGGCCAGTCCTAGCGTGGGTATCTATCGGAGTTAGGAAAGAGTGTGTGGTGGTGTCCAGCCTTGGTTCTCCTGCGTAATAGC<br>TGGCGTAGTAGCGCTCTGAGTTAGGAAAGTCTGCTCAGCGTACCGC |
| 10 / Left end | 72 | 59.7 | 93 | 16417.0 | 17126.0 | | GGGGTGGGCTGGGTGGGTAGCTCCTCTGTGTGGTCTTGAGCTCATCTCTTGCGGCGTAGGAAAGGTGGTTGT |
| 11 / Right end | 73 | 60.3 | 75 | 9173.0 | 9482.5 | | GGGCCCTTGCGAGGAGCTGGAAAGGGCTATTGTGGCAGCGTACGGCTGAGTGACTGCCTTTGTGTACTGCTGC |
| 11 / Left end | 152 | 41.4 | 82 | 3672.0 | 3772.8 | | GGGGAGCAATCTGAGGGCCTTTTTGATTCTATTGGGGAACAGCTAGGGATCAGGAAAGGCTGTGACAATTTATTCAGAATTGTTTGGTTAAG<br>GATTCACAGAGATATTTGGGTTGCAAAAGGTATGGTTTTGTGAATCTGTGGTAGTACGAT |
| 12 / Right end | 83 | 66.3 | 71 | 15769.0 | 16413.0 | | GGGGTGCGTTGGAGGGTGTGGCGAACTCTGGGGCGTCCGTAGCTTGCCGTGGGTCGCTCACAGGGCTGAGGAAAGAGGGAGCA |
| 12 / Left end | 153 | 50.3 | 74 | 4111.0 | 4250.8 | | GGGGGGTTATTGATTTTTGATCTCCGTAATCTGGCGCAGACCCCCAGAATCATCTTTGAACCACCGTATCTGGGTCAGGAAAGGGCGTATTG<br>CTGTGGGACCTGGTGCATCTTGTAGAGCTTGGGGAGAGCTTTCTATCGGTACCAAACACTT |
| 13 / Right end | 52 | 61.5 | 78 | 3423.5 | 3864.8 | (a) | GGGGTGGGTGTGGCTCTTTGGCAGCGTAGCTTGTAGTGCGAGGGCCAGGAAA |
| 13 / Left end | 73 | 58.9 | 89 | 4293.0 | 4419.0 | | GGGCCGAGGGAGTGCCTTTTTGGGTTTCCGTGGATCCGAGCGAGGAGCTGGAAAGGGCTCTTGTGTTAGCGTA |
| 14 / Right end | 72 | 65.3 | 112 | 3586.0 | 3715.0 | | GGGAAAGGGCGTACTGCTGCCTGTGGCCTGCGTGGTGGTCCCCAACGGTCCGTAGCGGATCTGGTGCGTAGT |
| 14 / Left end | 63 | 65.1 | 77 | 5597.0 | 5877.0 | | GGGCTCTTGGCTTTTGCGCCTGTCGAGGGTCAGGAACGGGCGATTGCGGCAGCGTAGTGGCGT |
| 15 / Right end | 72 | 52.8 | 102 | 1480.0 | 1588.3 | | GGGTGCTGGAAAGGGTGTAGTGTTGGTTTTGAGTGGCGTAGTTGGCTGATCCTGTGTCTGTGGTGACTACGA |
| 15 / Left end | 73 | 61.6 | 96 | 1641.5 | 1755.5 | | GGGAGGGCTCGGGAAAGAGGGCATAGGTCAGCGTACGGGTGGTGTACTGGCTTGCTCTGGGCCTGTGGTTACT |

| | | | | | | | |
|---|---|---|---|---|---|---|---|
| 16 / Right end | 59 | 57.6 | 139 | 20598.0 | 21583.0 | | GGGGGTGCGGCTGATTGGCTGTGTGTGGTGATCTGATTGATCTGGTGCGGCGTAGGAAA |
| 16 / Left end | 72 | 58.3 | 84 | 16277.0 | 17099.3 | | GGGGCTGGCGTTTGGTGTGATTGGCGTGTCCTTGTGTGAGTGGTGCGTAGTGCCCTCCTTTTGGTCTGGCAA |
| 17 / Right end | 72 | 59.7 | 94 | 2776.5 | 2942.5 | | GGGAGTGGGAAAGGGCCTGGTGTGGCTTTTGAGCGGCGTAGCTGGCTGGTCTCTGTTTGCTGGTGACTACCA |
| 17 / Left end | 151 | 49.7 | 97 | 5481.0 | 5595.0 | | GGGGTTACGGCGAGGGATCTTGGTGAGACTTGGTGGGTGTGTTGTGGTGGTGCTTGTGGCCTCAGAACCGCCTTTAACCGCACAATTTCTCT CTGAATTAGGAAAGATCTATTCTTAGCTACATCCAGCGTAGTATAGAAGGGTGTTCCCA |
| 18 / Right end | 178 | 60.1 | 114 | 18953.5 | 19670.5 | (b) | GGGGGATCAGTGCTAGCGTGGGTGCCCTCTAGGAATTAGGAAAGAGTGAGTGACGGATTGCGAGCTAGGGAAGGTATCTGGGCCTTTTGG TGCACAGCTTGGCGATCGAACCACCGGCGTAGCGTAGCCGGCGGTGGTGACTACCGGCGGATAGGCAAGTGGTGCGCTTGTGGCGTAC |
| 18 / Left end | 83 | 60.2 | 74 | 5558.5 | 5706.8 | | GGGCGTGAGGCGGCGTACGGGCATTGTCCTGTTCTGGAGACATCTCAGAGGGCTGAGGAAAGCCATGGTGTGTGCGTTACGGA |
| 19 / Right end | 164 | 54.3 | 74 | 6071.5 | 6228.5 | | GGGGGTGTCTGGCTCTGTTGTGGTGCCCTGTGTTGAGAGGTGTGTTCAGTGGTTGGCGGCGTAATGGGGTTGGTGTGGTCTACAGTGTGGC TGGTGTTGGCTTGACTTTTCTAGCAGACTCAGGAAAGCCAGAACAGCAAGGATAGAGTGGTGTACGGTGTTGT |
| 19 / Left end | 74 | 58.1 | 54 | 18197.0 | 19038.3 | | GGGTGGAGGGCTCGTGTTTTGGCCCCCGTGGGTGACTTTTCGAGGCACAGGAAAGCGAAGCTGGATGTCTAACT |
| 20 / Right end | 130 | 60.0 | 94 | 6351.5 | 7029.0 | (c) | GGGGGGTCCTCCTCTGACTTGGGAAAGAGTGCGTGATTGAGATCCGGTAGGGCGGAGTGCTGATTGGGGGTCCTCTTTTGGGGCGGGAAA GGGTGGGTGGGTGGTGTTCTTGCGTGGCTCAGTGCTTAGT |
| 20 / Left end | 130 | 60.0 | 106 | 3887.5 | 4178.0 | (c) | GGGGGGTCCTCCTCTGACTTGGGAAAGAGTGCGTGATTGAGATCCGGTAGGGCGGAGTGCTGATTGGGGGTCCTCTTTTGGGGCGGGAAA GGGTGGGTGGGTGGTGTTCTTGCGTGGCTCAGTGCTTAGT |

a, b or c - these motifs are identical

**Supplementary Table S3B.** Telomeric sequences of *J. pallidilutea*

| Chromosome / position | Size [bp] | G+C% | # reads | median [bp] | Q3 [bp] | Note | Motif [5'->3'] |
|---|---|---|---|---|---|---|---|
| **Proximal repeats** | | | | | | | |
| 2 / Left end | 56 | 46.4 | 58 | 916.5 | 1205.3 | (c) | GGGTACTTGAAGAACAGTGTTGGTCTTTATCTGTAGCTGTTGTCCGCTTCTTGCGA |
| 2 / Left end | 57 | 50.9 | 58 | 242.0 | 844.0 | | GGGTACTTGAAGAACAGTTGTTGGTCTCTATCCGTTGTCCTCCTCCGCTTCTTGCGA |
| 3 / Right end | 58 | 53.4 | 73 | 3282.0 | 3375.0 | (e) | GGGTACTTGAAGAACAGTTGTTGGTCTCTATCCGTCGTCCCTCCTCCGCTTCTTGCGA |
| 4 / Right end | 60 | 53.3 | 62 | 2387.0 | 2512.0 | | GGGGATCTGTGTTGCTCTGTCCGGATCTAGGGGCTCTGAAGTGCCTCTTATCTGATCTGT |
| 5 / Right end | 56 | 51.8 | 72 | 3206.5 | 3288.0 | | GGGTACTTGAAGAACAGTGTTGGTCTCTATCCGTTGTCCTCCTCCGCTTCTTGCGA |
| 6 / Right end | 57 | 47.4 | 77 | 443.0 | 481.0 | (d) | GGGTACTTGAAGAACAGTTGTTGGTCTTTATCTGTAGCTCTTGTCCGCCTCTTGCGA |
| 7 / Right end | 43 | 55.8 | 64 | 257.0 | 310.8 | | GGGGTGTTGTTGTCCTTGTCCGCTGCTTGCGAGGGTACTTGAA |
| 9 / Left end | 58 | 53.4 | 56 | 998.5 | 1017.3 | (e) | GGGTACTTGAAGAACAGTTGTTGGTCTCTATCCGTCGTCCCTCCTCCGCTTCTTGCGA |
| 10 / Right end | 56 | 51.8 | 70 | 180.0 | 198.0 | | GGGTACTTGAAGAACCGTTGTATCGTCTTAGTCATCGCCCTGTCCGCTTCTTGCGA |
| 10 / Left end | 57 | 47.4 | 66 | 1684.5 | 1927.5 | (d) | GGGTACTTGAAGAACAGTTGTTGGTCTTTATCTGTAGCTCTTGTCCGCCTCTTGCGA |
| 12 / Right end | 58 | 53.4 | 89 | 4615.0 | 4813.0 | (e) | GGGTACTTGAAGAACAGTTGTTGGTCTCTATCCGTCGTCCCTCCTCCGCTTCTTGCGA |
| 13 / Right end | 56 | 46.4 | 64 | 221.5 | 234.8 | (c) | GGGTACTTGAAGAACAGTGTTGGTCTTTATCTGTAGCTGTTGTCCGCTTCTTGCGA |
| 14 / Right end | 56 | 48.2 | 65 | 192.0 | 238.0 | | GGGTACTTGAAGAACCGTTGTATTGTCTTAGTCATAGCCTCCTCCGCTTCTTGCGA |
| 14 / Right end | 96 | 47.9 | 65 | 489.0 | 509.0 | | GGGTACTTGAAGAACCGTTCGTTTCTGTTCGGTCTCTGTCGAGGGTACATGAAGAGTGTTGTCATTCCTTTTCATAGTCCTTTTCGCCCCTTGCGA |
| 14 / Right end | 58 | 53.4 | 65 | 1032.0 | 1108.0 | (e) | GGGTACTTGAAGAACAGTTGTTGGTCTCTATCCGTCGTCCCTCCTCCGCTTCTTGCGA |
| 14 / Right end | 99 | 51.5 | 65 | 1167.0 | 1217.0 | | GGGGTGTTGTTGTCCTCCTCCGCTTCTTGCGAGGGTACTTGAAGAACCGTTGTATTGTCTTAGTCATAGCCTCCTCCGCTTCTTGCGAGGGTACTTGAA |
| 15 / Left end | 101 | 46.5 | 56 | 640.5 | 883.0 | | GGGTACTTGAAGAACCATTCGTTTCTGTTCGGTATCTGTCGAGGGTACATGAAGAGTGTTGTCATTCATTGTCATTCATAGTCCCTGTCCGCCTCTTGCGA |
| **Distal repeats** | | | | | | | |
| 1 / Right end | 81 | 38.3 | 66 | 5794.0 | 5899.0 | (f) | GGGTAAATGAAGGTATATCTATTTCCTTAGTCTATTTTCGATTGATTCCAAGGTCTCTGTGTCCATTGCAATGGGGATCTA |
| 1 / Left end | 96 | 44.8 | 53 | 2239.0 | 2760.0 | | GGGTACTTGAAGAACTATCTGTGCTGTTTGGTTGTCTGTCTAGGGTACATGAAGAGTGTTGTGATTCATTGTCATTGTCCTTTTCGCCCCTTGCGA |
| 2 / Right end | 63 | 44.4 | 64 | 3759.5 | 3880.5 | | GGGTAAATGAAGGGTTGTCATTGGTCCTCGTCGCCATTGTCTCTGTTTTGTAATCGGTTTCTA |
| 2 / Left end | 100 | 46.0 | 58 | 1042.0 | 2938.5 | (h) | GGGTACTTGAAGAACTATCTGTGCTGTTTGGTTGTCTGTCTAGGGTACATGAAGAGTGTTGTGATTCATTGTCATTCATTGTCCTGTCCGCCCCTTGCGA |
| 3 / Right end | 100 | 46.0 | 73 | 2064.0 | 2399.0 | (h) | GGGTACTTGAAGAACTATCTGTGCTGTTTGGTTGTCTGTCTAGGGTACATGAAGAGTGTTGTGATTCATTGTCATTCATTGTCCTGTCCGCCCCTTGCGA |
| 3 / Left end | 41 | 51.2 | 81 | 6763.0 | 7912.0 | (a) | GGGTGCTTGAAGAGCGTTGTCTGTCTGTGGTTGTAGTGTGA |
| 4 / Right end | 81 | 38.3 | 62 | 2295.0 | 2453.0 | (f) | GGGTAAATGAAGGTATATCTATTTCCTTAGTCTATTTTCGATTGATTCCAAGGTCTCTGTGTCCATTGCAATGGGGATCTA |
| 4 / Left end | 96 | 46.9 | 59 | 3052.0 | 3631.5 | (g) | GGGTACTTGAAGAACTATCTGTGCTGTTTGGTTGTCTGTCTAGGGTACATGAAGAGTGTTGTGATTCATTGTCATTGTCCTGTCCGCCCCTTGCGA |
| 5 / Right end | 100 | 46.0 | 72 | 3351.0 | 4267.5 | (h) | GGGTACTTGAAGAACTATCTGTGCTGTTTGGTTGTCTGTCTAGGGTACATGAAGAGTGTTGTGATTCATTGTCATTCATTGTCCTGTCCGCCCCTTGCGA |
| 5 / Left end | 55 | 49.1 | 90 | 1641.0 | 3786.8 | | GGGAGTGTTTTGGTAGTGCCTGTGCGAGGCTATTTGAAGCCACTTGTGATCCTTT |
| 6 / Right end | 100 | 46.0 | 77 | 4066.0 | 5331.0 | (h) | GGGTACTTGAAGAACTATCTGTGCTGTTTGGTTGTCTGTCTAGGGTACATGAAGAGTGTTGTGATTCATTGTCATTCATTGTCCTGTCCGCCCCTTGCGA |
| 6 / Left end | 42 | 54.8 | 76 | 1576.5 | 3210.0 | (b) | GGGTACGTGAAGCGTAGTCCGTGTCTGTTCAGTCTCTGTCGA |
| 7 / Right end | 100 | 46.0 | 64 | 4074.5 | 5419.5 | (h) | GGGTACTTGAAGAACTATCTGTGCTGTTTGGTTGTCTGTCTAGGGTACATGAAGAGTGTTGTGATTCATTGTCATTCATTGTCCTGTCCGCCCCTTGCGA |
| 7 / Left end | 96 | 46.9 | 67 | 3111.0 | 4956.5 | | GGGTACTTGAAGAACCGTTCGTTTCTGTTCGGTCTCTGTCGAGGGTACATGAAGAGTGTTGTCATTCCTTTTCATAGTCCTTTTCGCTGCTTGCGA |
| 8 / Right end | 96 | 46.9 | 63 | 3182.0 | 3717.5 | (g) | GGGTACTTGAAGAACTATCTGTGCTGTTTGGTTGTCTGTCTAGGGTACATGAAGAGTGTTGTGATTCATTGTCATTGTCCTGTCCGCCCCTTGCGA |
| 8 / Left end | 62 | 41.9 | 67 | 4461.0 | 5001.5 | | GGGGACTGAGGGTAAATGAAGCATATTTCAGAGCATGCAAAGGCAATAGAGTCGATCTAAAA |

| | | | | | | | |
|---|---|---|---|---|---|---|---|
| 9 / Right end | 84 | 52.4 | 72 | 4665.0 | 6432.8 | | GGGGTTGCTCCCTGTCCATCCAGTCACTTGCTAGGAGATCTGAAGGGGTTTGTCCTTGTCTTTTCCTGTTTTCCCAGTGCCGTA |
| 9 / Left end | 100 | 46.0 | 56 | 1986.0 | 2389.5 | (h) | GGGTACTTGAAGAACTATCTGTGCTGTTTGGTTGTCTGTCTAGGGTACATGAAGAGTGTTGTGATTCATTGTCATTCATTGTCCTGTCCGCCCCTTGCGA |
| 10 / Right end | 100 | 46.0 | 70 | 4288.5 | 5191.3 | (h) | GGGTACTTGAAGAACTATCTGTGCTGTTTGGTTGTCTGTCTAGGGTACATGAAGAGTGTTGTGATTCATTGTCATTCATTGTCCTGTCCGCCCCTTGCGA |
| 10 / Left end | 96 | 45.8 | 66 | 1056.0 | 1883.8 | | GGGTACTTGAAGAACTATCTGTGCTGTTTGGTTGTCTGTCTAGGGTACATGAAGAGTGTTGTGATTCATTGTCATTGTCCTGTCCGCCCTTTGCGA |
| 11 / Right end | 83 | 42.2 | 87 | 6907.0 | 7815.5 | | GGGGTGTGAGGGTCTCTGAAGCCATAAAGGGTGTTCTTTGGTGCTTGTTCCTCGTTTTATTGTTGTTTTTGTGTGTTTTCATT |
| 11 / Left end | 42 | 54.8 | 133 | 1780.0 | 2233.0 | (b) | GGGTACGTGAAGCGTAGTCCGTGTCTGTTCAGTCTCTGTCGA |
| 12 / Right end | 99 | 45.5 | 89 | 1352.0 | 1926.0 | | GGGTACTTGAAGAACTATCTGTGCTGTTTGGTTGTCTGTCTAGGGTACATGAAGAGTGTTGTGATTCATTGTCATTCATTGTCCTGTCCGCCCTTGCGA |
| 12 / Left end | 75 | 38.7 | 80 | 2181.5 | 2374.5 | | GGGTAAATGAAGGGTGTTTGTAGGTTGTTAATTGGCCCCTTAACGGTTTTTGATTGGTTTTGTAATCGGTTTCTA |
| 13 / Right end | 96 | 46.9 | 64 | 3485.5 | 3846.8 | (g) | GGGTACTTGAAGAACTATCTGTGCTGTTTGGTTGTCTGTCTAGGGTACATGAAGAGTGTTGTGATTCATTGTCATTGTCCTGTCCGCCCCTTGCGA |
| 13 / Left end | 81 | 40.7 | 56 | 1394.5 | 1481.8 | | GGGATCTAGGGTAAATGAAGGTATATGGTCATCCTTAGGCGATTTTTGCTTAATCCAGATGTGTCTGTATGGCCATAGATA |
| 14 / Right end | 100 | 46.0 | 65 | 2115.0 | 2339.0 | (h) | GGGTACTTGAAGAACTATCTGTGCTGTTTGGTTGTCTGTCTAGGGTACATGAAGAGTGTTGTGATTCATTGTCATTCATTGTCCTGTCCGCCCCTTGCGA |
| 14 / Left end | 41 | 51.2 | 72 | 5615.5 | 6880.5 | (a) | GGGTGCTTGAAGAGCGTTGTCTGTCTGTGGTTGTAGTGTGA |
| 15 / Right end | 81 | 46.9 | 67 | 4819.0 | 5073.5 | | GGGATTGAGGGTAAATGAAGGTATCTGTGTTGCCTTTGGTCATTCTGGGTTCATCTGGAGGTGCCTGTATCGCCATAGATA |
| 15 / Left end | 100 | 46.0 | 56 | 3293.5 | 3777.5 | (h) | GGGTACTTGAAGAACTATCTGTGCTGTTTGGTTGTCTGTCTAGGGTACATGAAGAGTGTTGTGATTCATTGTCATTCATTGTCCTGTCCGCCCCTTGCGA |
| 16 / Right end | 41 | 51.2 | 64 | 4443.5 | 4695.8 | (a) | GGGTGCTTGAAGAGCGTTGTCTGTCTGTGGTTGTAGTGTGA |
| 16 / Left end | 46 | 50.0 | 50 | 1456.0 | 3156.8 | | GGGTACCTGAAGAGTGTTGTCTGTCCTTCCCTTTGTTCTCTGTCGA |

a, b, c, d, e, f, g or h - these motifs are identical

**Supplementary Table S3C.** Telomeric sequences of *P. phylloscopi*

| Chromosome / position | Size [bp] | G+C% | # reads | median [bp] | Q3 [bp] | Note | Motif [5'->3'] |
|---|---|---|---|---|---|---|---|
| **Proximal repeats** | | | | | | | |
| 2 / Right end | 101 | 58.4 | 115 | 1509.0 | 1523.0 | | GGGTTTGAAATGACCATTGCCATCCGTTGTGGCGCTGTAGAGTGCCTTCTGAGTGGCGTGGCGCCTTTCCCTGCCTTCCCCTGTCTGCCAACGCGTTGTGA |
| 3 / Left end | 81 | 63.0 | 114 | 3106.5 | 3184.0 | | GGGGCTGGGAGGGTCGTGTGTCTTCCGTCTTAGCCTGTGCTGGCGTGAGGGGCTGGAATGGGTGGCGGTGTTATAGTGCGA |
| **Distal repeats** | | | | | | | |
| 1 / Right end | 132 | 61.4 | 101 | 9174.0 | 9833.0 | | GGGCACCTTCCGAGGCCTTAGGAGTGACAGGGATCGGATTGCTATTGCTGCCAGAGTGTGTGCCGGGCTGCACCTGCCCTCCCCACCGAGATAGGCGTCGTGAGGATTGGAAAAGGGTCTTCCAGTCCCCCA |
| 1 / Left end | 121 | 52.9 | 91 | 8075.0 | 8618.0 | | GGGGTACCTTCTAGGTGCTTAGGAGTGACAACGATCCGATTGTTGTTGCGCACGTGGTGTGCCACTACCCGTACTGTGCCCGGTGTCAGAGTGAGGATTTGAAAAGCGTCAATACAGCTGT |
| 2 / Right end | 130 | 54.6 | 115 | 15550.0 | 16655.5 | (a) | GGGGCTGGAACGGGTAGCGGTGTCAGTGAGGGTTTGGGAGGGCCGTATGGTATCTGCCAACGCGTTGTGAGGGTTTGAAATGCCATATAGTATCGCCGTGGTATAGTGTGAGGGTTAGCGATGAGTGTGA |
| 2 / Left end | 101 | 59.4 | 111 | 5939.0 | 6439.0 | | GGGGTTGGAAAGTGCATTGCTGTCCGTTGTGGCACTGTAGAGTGCGTTTGGAGGGCGGTAGCGCCTTTCTCTGTCCTCTGGTGTCCGTGGCAGCGCTGTGA |
| 3 / Right end | 101 | 53.5 | 77 | 17727.0 | 19171.0 | | GGTCATCGGTGGCATCCGTTGTCGCAGTTGTAGAGTGAGGATCAGCAATGAGTGGTGTGGTGTCGTCTGTGGCACCTGGTGATAGAGTGAGGATCAGCAAT |
| 3 / Left end | 130 | 54.6 | 114 | 3584.0 | 4082.0 | (a) | GGGGCTGGAACGGGTAGCGGTGTCAGTGAGGGTTTGGGAGGGCCGTATGGTATCTGCCAACGCGTTGTGAGGGTTTGAAATGCCATATAGTATCGCCGTGGTATAGTGTGAGGGTTAGCGATGAGTGTGA |

a - these motifs are identical

**Supplementary Table S3D.** Telomeric sequences of *S. kandeliae*

| Chromosome / position | Size [bp] | G+C% | # reads | median [bp] | Q3 [bp] | Note | Motif [5'->3'] |
|---|---|---|---|---|---|---|---|
| **Proximal repeats** | | | | | | | |
| 1 / Left end | 69 | 49.3 | 54 | 3790.5 | 3963.5 | | GGGATCTACCACTGGTGTTGTGTTGTCCTGGCTGTTGTCGTGTCGGTGTTTAGTTTTGACTAAGGCGTA |
| 2 / Right end | 69 | 52.2 | 49 | 4366.0 | 4489.0 | | GGGATCCACGACTGGTGTTGTGTTGCCGTGCCTGTTGTCGTGTGCTGCTTTTGTTTTGACTAAGGCGTA |
| 2 / Right end | 62 | 51.6 | 49 | 8061.0 | 9508.0 | | GGGACTGGTGTTGTGTTGCCGTGCCTGTTGTCGTGTGCTGCTTTTGTTTTGACTAAGGCGTA |
| 2 / Left end | 111 | 57.7 | 84 | 6941.0 | 7881.3 | | GGGGCCGTTGGCCCGTGTGAGTTCCCTGTTGTTGTTAGCCCTTCGCTATGACTGAGGGGAAGGGATCAATGGGTAGCCTAGTTGCCTGTTTA GTTTTGGCGCCGTTGGCGC |
| **Distal repeats** | | | | | | | |
| 1 / Right end | 70 | 57.1 | 58 | 18862.0 | 21709.5 | (a) | GGGTTGGGTCCTGTGTCGAGGGTTGGGTATCCGTGCCTTGTCTCTGTGGTTGTCTGTGGTGCTTGTCTGT |
| 1 / Left end | 62 | 48.4 | 54 | 17025.5 | 26422.5 | (b) | GGGACTGGTGTTGTGTTGTCCTGGCTGTTGTCGTGTCGGTGTTTAGTTTTGACTAAGGTGTA |
| 2 / Right end | 62 | 48.4 | 49 | 3688.0 | 9123.0 | (b) | GGGACTGGTGTTGTGTTGTCCTGGCTGTTGTCGTGTCGGTGTTTAGTTTTGACTAAGGTGTA |
| 2 / Left end | 111 | 57.7 | 84 | 10572.5 | 13213.3 | | GGGGCCGTTGGCCCGTGTGAGTTCCCTGTTGTTGTTAGCCCTTCGCTATGACTGAGGGGAAGGGATCCATGTGTAGGCTAGTTGCCTGTTTA GTTTTGGCGCCGTTGGCGC |
| 3 / Right end | 62 | 48.4 | 31 | 14004.0 | 17434.5 | (b) | GGGACTGGTGTTGTGTTGTCCTGGCTGTTGTCGTGTCGGTGTTTAGTTTTGACTAAGGTGTA |
| 3 / Left end | 70 | 57.1 | 50 | 11772.0 | 18335.8 | (a) | GGGTTGGGTCCTGTGTCGAGGGTTGGGTATCCGTGCCTTGTCTCTGTGGTTGTCTGTGGTGCTTGTCTGT |
| 4 / Right end | 61 | 55.7 | 73 | 20821.0 | 24882.0 | | GGTGTCTGGTGCGGTTGCGGTTCCGAGGCTTAGGTATCCGTGTGTTGTCTCTGTTGTGTGT |
| 4 / Left end | 62 | 50.0 | 43 | 14728.0 | 20394.0 | | GGGACTGGTGTTGTGTTGTCCTGGCTGTTGTCGTGTCGGTGTTTAGTTTTGACTAAGGCGTA |
| 5 / Right end | 70 | 57.1 | 46 | 11721.5 | 23547.8 | (a) | GGGTTGGGTCCTGTGTCGAGGGTTGGGTATCCGTGCCTTGTCTCTGTGGTTGTCTGTGGTGCTTGTCTGT |
| 5 / Left end | 122 | 54.9 | 33 | 15422.0 | 23408.0 | | GGGGCAGGTATCCGTTGTTGTTACTTAGTGGTCGTTTGTTTTAGGGCGAGGCGTAGGGATCTACCACTGGGTGGCGTTGTGATTGACCGGT GGTGTGCGTGTCTGTCTTGGTTCGGTTGCGA |
| 6 / Right end | 55 | 52.7 | 47 | 10005.0 | 20887.0 | (c) | GGGGTAGGGATCCGTTGGTGTTCCTTCTTCTTCTGCTCCTGTTGTGTTGTGCTGA |
| 6 / Left end | 55 | 52.7 | 40 | 9679.0 | 18302.0 | (c) | GGGGTAGGGATCCGTTGGTGTTCCTTCTTCTTCTGCTCCTGTTGTGTTGTGCTGA |

a, b or c - these motifs are identical

**Supplementary Table S4. List of hybridization probes.** The oligonucleotides derived from distal telomeric repeats were synthesized by Microsynth AG.

| Oligonucleotide name | Chromosome | Telomere | Strand | Sequence [5'->3'] |
|---|---|---|---|---|
| motif01s | 1 | Left | G-rich | GGCTGGCTTTTGTTGTGAGCGGCGTGGGTGCGTGGCACTGGTCCGTCGAGTTCTCATTTGGGTCTGGTAAGG |
| motif01e | 1 | Right | G-rich | GTAGGGGGCTGCGTGGCCTCGTGTGCTGCGTGTGGCTACGGTAGCCGATCTCATCTGACTGAGGAAAGAGGGC |
| motif03s | 3 | Left | G-rich | TACCTGGGCTGCTGTAGCCGTAACTGCTCTGTCAAGGCACAGGAAAGGGTCATGGTATGGTTTGCTTGTGGCG |
| motif03e | 3 | Right | G-rich | ACGACCTTGCTGTGCGCCGTTGCTTTGCTCTGGTGGATCAGGAAAGATATGGTGTGTGGTTTTGAGTGGCGT |
| motif15s | 15 | Left | G-rich | GTACGGGTGGTGTACTGGCTTGCTCTGGGCCTGTGGTTACTGGGAGGGCTCGGGAAAGAGGGCATAGGTCAGC |
| motif15s_anti | 15 | Left | C-rich | GCTGACCTATGCCCTCTTTCCCGAGCCCTCCCAGTAACCACAGGCCCAGAGCAAGCCAGTACACCACCCGTAC |
| motif15e | 15 | Right | G-rich | GGGTGTAGTGTTGGTTTTGAGTGGCGTAGTTGGCTGATCCTGTGTCTGTGGTGACTACGAGGGTGCTGGAAA |
| motif15e_anti | 15 | Right | C-rich | TTTCCAGCACCCTCGTAGTCACCACAGACACAGGATCAGCCAACTACGCCACTCAAAACCAACACTACACCC |
| motif20se | 20 | Both | G-rich | GGGTGGGTGGGTGGTGTTCTTGCGTGGCTCAGTGCTTAGTGGGGGGTCCTCCTCTGACTTGGGAAAGAGT |

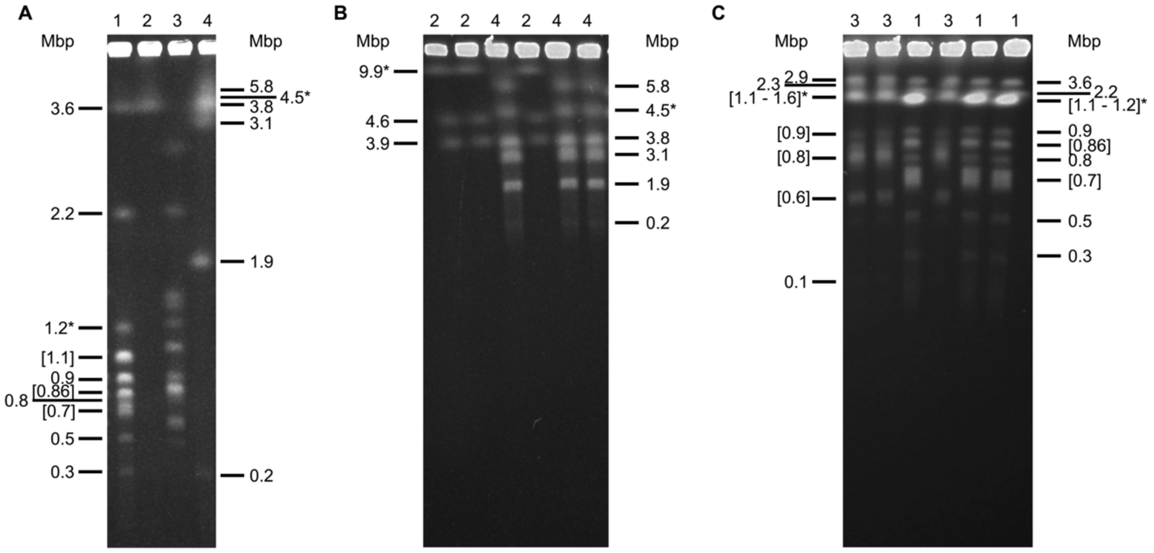


**Supplementary Figure S1. Electrophoretic karyotypes analyzed using PFGE.** DNA samples of *J. pallidilutea* (lane 1), *P. phylloscopi* (lane 2), *J. angkorensis* (lane 3), and *S. kandeliae* (lane 4) were prepared in agarose plugs and electrophoretically separated (see Supplementary Methods for details). The separations were performed in 0.5×TBE at 9 °C and following settings: (A) a 0.8% (w/v) agarose gel; pulses from 60 to 600 s (linear interpolation) for 72 hours at 100 V; (B) a 0.8% (w/v) agarose gel; pulses from 800 to 3600 s (linear interpolation) for 96 hours at 54 V; and (C) a 0.9% (w/v) agarose gel; pulses from 60 to 135 s (linear interpolation) for 26 hours at 145 V. The sizes of bands containing multiple co-migrating chromosomes are shown in square brackets; an asterisk indicates the chromosome containing the rDNA cluster arrays in the corresponding species.

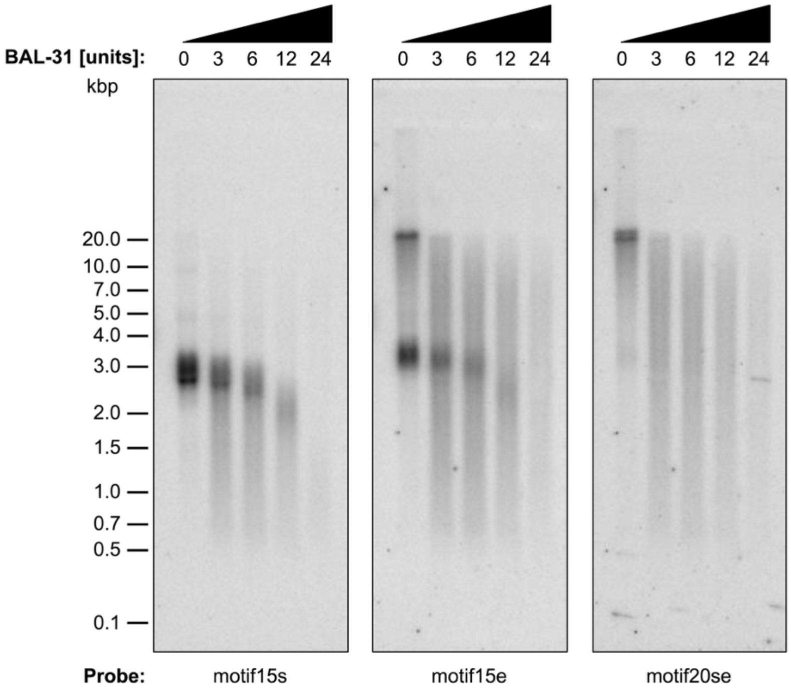


**Supplementary Figure S2. Noncanonical telomeric arrays are sensitive to BAL-31 nuclease.** Total cellular DNA of *J. angkorensis* was first digested with 0, 3, 6, 12, and 24 units of BAL-31 nuclease and then cleaved by restriction endonuclease *Eco*RV. The resulting DNA fragments were electrophoretically separated in a 0.7% (w/v) agarose gel, blotted onto a nylon membrane, and hybridized with radioactively labeled oligonucleotide probes derived from telomeric repeats of chromosome 15 (motif15s, motif15e) and chromosome 20 (motif20se) (**Supplementary Table S4**; see Supplementary Methods for details).

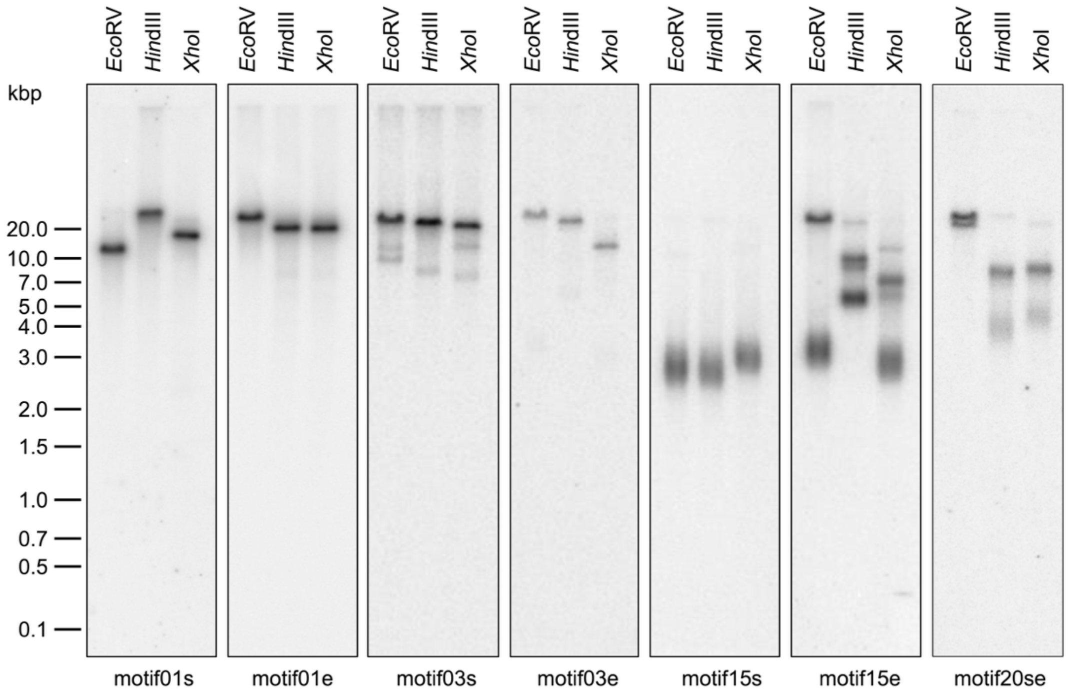


**Supplementary Figure S3. The lengths of *J. angkorensis* telomeres analyzed using TRF assay.** Total cellular DNA of *J. angkorensis* was digested with indicated restriction endonucleases. The resulting DNA fragments were electrophoretically separated in a 0.7% (w/v) agarose gel, blotted onto a nylon membrane, and hybridized with radioactively labeled oligonucleotide probes derived from the telomeric repeats of chromosome 1 (motif01s, motif01e), chromosome 3 (motif03e, motif03s), chromosome 15 (motif15s, motif15e), and chromosome 20 (motif20se) (**Supplementary Table S4**; see Supplementary Methods for details).

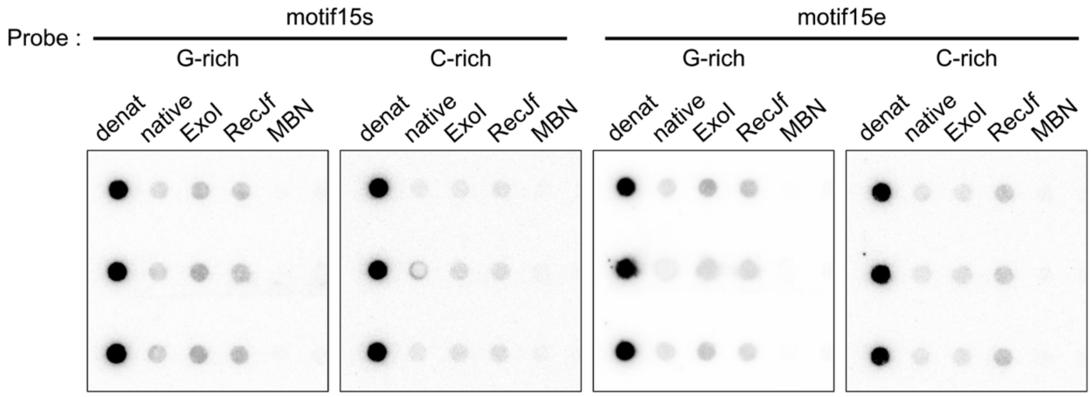


**Supplementary Figure S4. Telomeric arrays contain regions with single-stranded DNA.** Samples containing denatured, native control and nuclease-digested DNA samples of *J. angkorensis* were analyzed by dot blot hybridization with radioactively labeled oligonucleotide probes derived from telomeric repeats of chromosome 15. G-rich probes: motif15s, motif15e; C-rich probes: motif15s_anti, motif15e_anti (**Supplementary Table S4**; see Supplementary Methods for details). The experiment was performed in three technical replicates.

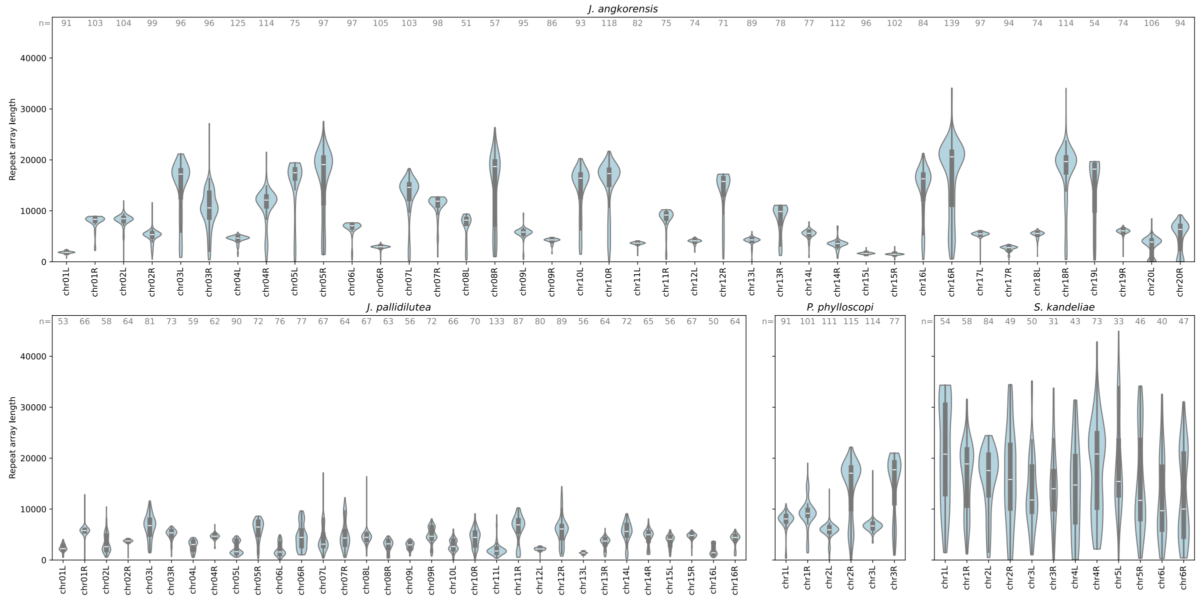


**Supplementary Figure S5. Violin plots of the lengths of telomere motif arrays in individual nanopore sequencing reads for each chromosomal end in all four studied species**. Both distal and proximal motifs are considered in the length. Only reads spanning a unique sequence region closest to chromosome end and longer than a species-specific threshold 20-30 kbp were considered. Nonetheless some reads may represent fragments not reaching the actual chromosome length. The numbers at the top of each plot show the read count used in each violin plot.

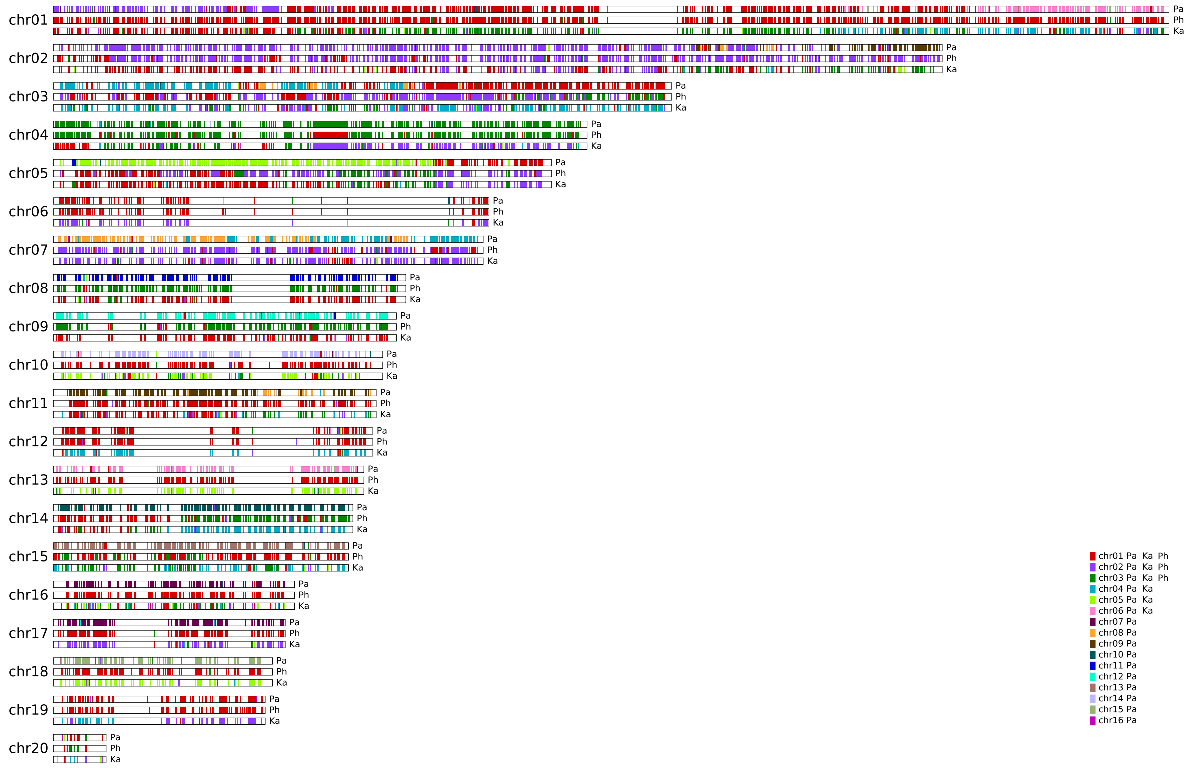


**Supplementary Figure S6. Chromosomes of *J. angkorensis* painted by nucleotide alignments with the three target genomes***.* Each *J. angkorensis* chromosome is shown three times, each time colored by a different target genome. Target genome is labeled near the right end of the chromosome using the first two letters of the species name (i.e. Pa for *J. pallidilutea,* Ph for *P. phylloscopi*, Ka for *S. kandeliae).* The color is based on the chromosome number in the target genome. Note that *J. pallidilutea*, *P. phylloscopi*, and *S. kandeliae* have 16, 3, and 6 nuclear chromosomes, respectively.

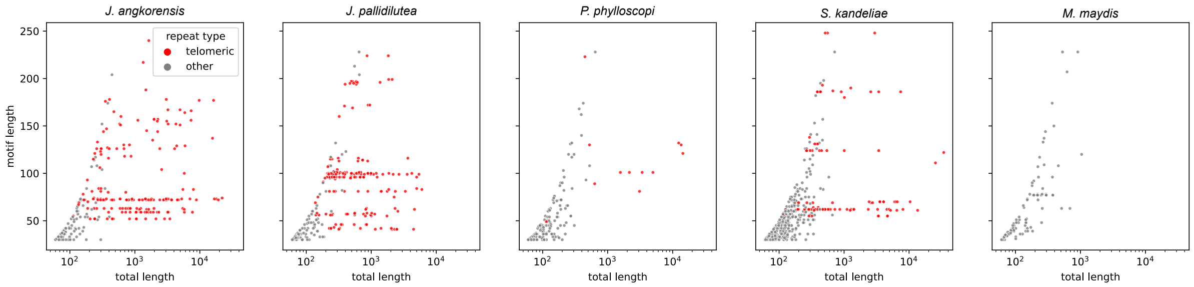


**Supplementary Figure S7. Tandem repeats present in the Microstromatales genomes**. All tandem repeats of length 30 to 250 bp with at least 2 full copies of the motif found by Tantan in the Microstromatales genomes. Each point represents one tandem repeat motif, with the total tandem array length as the x coordinate (in log scale) and the motif length as the y coordinate. Color distinguishes the repeats overlapping annotated telomeric arrays from others. In each species, the longest arrays are formed by telomeric motifs. Note that some found tandem repeats overlapping annotated telomeres are relatively short; these are often fragments separated by errors in sequence assembly or shorter stretches of proximal repeats. *M. maydis* is shown for comparison; its telomeric arrays have motif length (i.e. TTAGGG) below the 30 bp threshold.

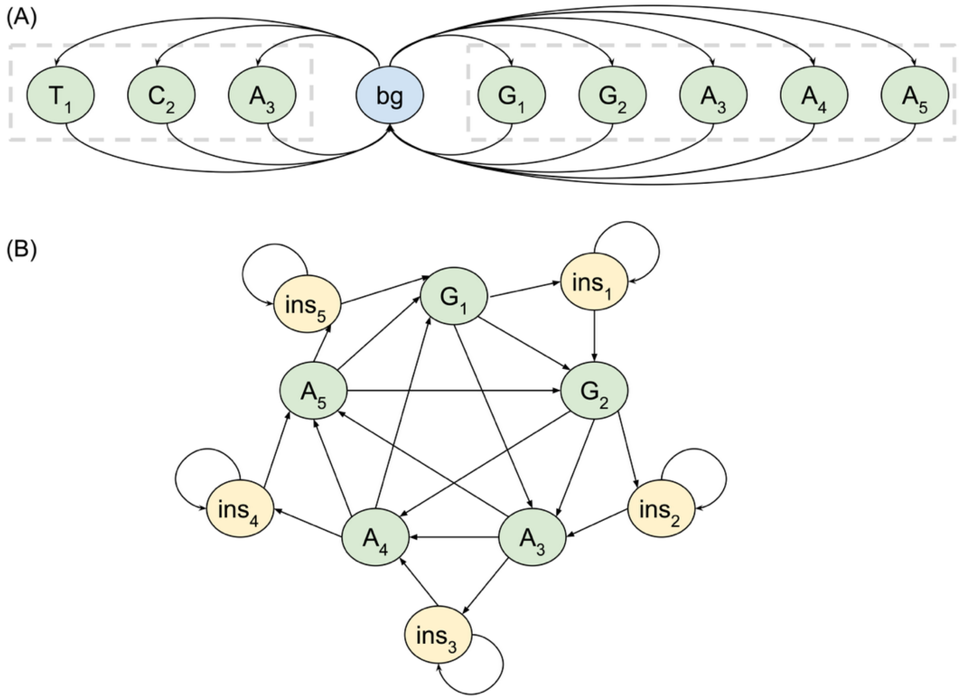


**Supplementary Figure S8. States of the HMM for finding telomeric motifs.** (A) Connections to/from the background state for a hypothetical chromosome end with motifs GGAA and TCA (in our application the motifs are much longer). (B) Connections within profile HMM for motif GGAAA. After each nucleotide, there is a separate insertion state. Deletions are handled by direct transitions between distant states. Here for simplicity at most one nucleotide can be deleted between two match states but in the actual HMM up to 7 nucleotides can be deleted, leading to 7 incoming and outgoing deletion transitions for each state.

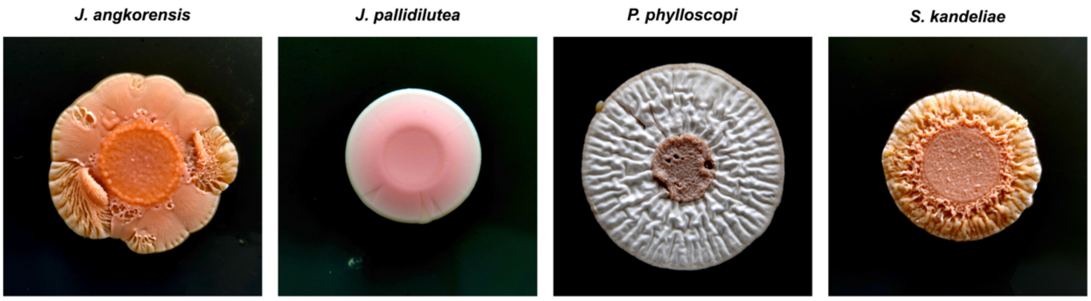


**Supplementary Figure S9. Colony morphology of the Microstromatales species.** The yeast cultures were grown on YM plates at 28 °C for 29 days.
